# Harmonized cross-species cell atlases of trigeminal and dorsal root ganglia

**DOI:** 10.1101/2023.07.04.547740

**Authors:** Shamsuddin A. Bhuiyan, Mengyi Xu, Lite Yang, Evangelia Semizoglou, Parth Bhatia, Katerina I. Pantaleo, Ivan Tochitsky, Aakanksha Jain, Burcu Erdogan, Steven Blair, Victor Cat, Juliet M. Mwirigi, Ishwarya Sankaranarayanan, Diana Tavares-Ferreira, Ursula Green, Lisa A. McIlvried, Bryan A. Copits, Zachariah Bertels, John S. Del Rosario, Allie J. Widman, Richard A. Slivicki, Jiwon Yi, Clifford J. Woolf, Jochen K. Lennerz, Jessica L. Whited, Theodore J. Price, Robert W. Gereau, William Renthal

## Abstract

Peripheral sensory neurons in the dorsal root ganglion (DRG) and trigeminal ganglion (TG) are specialized to detect and transduce diverse environmental stimuli including touch, temperature, and pain to the central nervous system. Recent advances in single-cell RNA-sequencing (scRNA-seq) have provided new insights into the diversity of sensory ganglia cell types in rodents, non-human primates, and humans, but it remains difficult to compare transcriptomically defined cell types across studies and species. Here, we built cross-species harmonized atlases of DRG and TG cell types that describe 18 neuronal and 11 non-neuronal cell types across 6 species and 19 studies. We then demonstrate the utility of this harmonized reference atlas by using it to annotate newly profiled DRG nuclei/cells from both human and the highly regenerative axolotl. We observe that the transcriptomic profiles of sensory neuron subtypes are broadly similar across vertebrates, but the expression of functionally important neuropeptides and channels can vary notably. The new resources and data presented here can guide future studies in comparative transcriptomics, simplify cell type nomenclature differences across studies, and help prioritize targets for future pain therapy development.

## Introduction

Despite the wide use of rodent chronic pain models to identify novel pain therapeutic targets, our understanding of the similarity between human and mouse somatosensory cell types remains limited. Mammals sense a range of external somatosensory stimuli through functionally distinct peripheral sensory neurons whose cell bodies reside in the trigeminal ganglion (TG) or the dorsal root ganglion (DRG)^1–4^. Improved insight into these neuronal subtypes and their molecular features in humans could lead to new therapeutic approaches for sensory nervous system disorders such as chronic pain and itch.

DRGs and TGs contain both neuronal and non-neuronal cell types. Non-neuronal cells such as satellite glia, Schwann cells, and fibroblasts, play important roles in ganglion structure as well as in modulating the electrophysiological properties of peripheral sensory neurons^5–7^. Peripheral sensory neurons are highly heterogenous and have been historically classified by combinations of features including their size, degree of myelination, nerve conduction velocity, environmental stimuli that evoke action potentials, and their gene expression. Large diameter, fast-conducting (2-55 m/s) neurons are termed A-fibers, and unmyelinated, slow conducting (< 2 m/s) neurons are termed C-fibers^8,9^. A-fibers tend to be specialized for but not exclusively sensitive to mechanical stimuli whereas C-fibers can detect a broader range of environmental stimuli including thermal, mechanical, and chemical stimuli^10,11^. While most of our knowledge of peripheral sensory neuron structure and function is related to their cutaneous innervation, it is important to note that peripheral sensory neurons also innervate a wider range of internal organs^12–14^.

The structure and function of distinct peripheral sensory neurons subtypes ultimately depends upon the genes they express, and much work has been dedicated to characterizing the molecular features that define each distinct subtypes^15,16^. Recent advances in single cell/nuclei RNA-seq (sc/snRNA-seq) have enabled unprecedented molecular insight into mammalian sensory neuron subtypes^11,17–24,24–26,26–32^. However, interpreting the conclusions between datasets remains challenging. Technical differences between studies, such as whole cell vs. nuclei preparation, sequencing platform, sequencing depth, and bioinformatic pipeline, can affect the number of distinct transcriptomic subtypes and the transcriptomic coverage per cell/nucleus^33,34^. Moreover, there are significant differences in nomenclature used to annotate transcriptomically-defined cells/nuclei across studies, which creates an additional challenge in interpreting the data between studies and species. Together, these challenges limit our ability to connect molecularly defined cell types with related structural and functional data, and contribute to uncertainty about the conservation of DRG and TG cell types across species^24,25,31,35^.

To address these challenges, we constructed harmonized cell atlases using sc/snRNA-seq data obtained from 15 DRG and 5 TG mammalian studies. These new reference atlases enabled the harmonization of cell type nomenclatures across studies, direct comparisons of cell types between studies and species, improved transcriptomic coverage of lowly expressed genes, and identification of rare cell populations within the sensory ganglia, including new immune subtypes. We then sequenced 135,834 nuclei from human DRG across three independent laboratories and found that the harmonized atlas significantly improved cell type annotations compared to when studies were annotated separately. Finally, we extended our molecular understanding of vertebrate sensory neuron evolution by performing scRNA-seq on axolotl DRGs and observed notable conservation of peripheral sensory neuron subtypes across broad classes of vertebrates. The harmonized somatosensory cell atlases (https://harmonized.painseq.com/) presented here provide a powerful new resource for integrating and annotating cell types in single-cell/nucleus transcriptomic/epigenomic datasets, comparing somatosensory cell types between species, and prioritizing targets for future pain therapeutic design.

## Results

### Harmonized mammalian somatosensory neuronal cell atlases

To construct the harmonized dorsal root ganglion (DRG) and trigeminal ganglion (TG) cell atlases, we developed a pipeline that minimizes study-specific batch effects (see Methods). Our approach includes a cell or nucleus in the final reference atlas only if its cell type annotation is consistent across two distinct computational pipelines, one that includes Seurat and one that includes LIGER (see Methods)^33,36,37^. A similar approach was used to build a harmonized spinal cord cell atlas ^38^.

We began constructing the harmonized reference atlas by first separating neurons and non-neurons from 316,897 DRG cells/nuclei across 15 studies (133,068 human, 15,892 rat, 100,607 mouse, 59,900 cynomolgus macaque and 7,430 rhesus macaque, 46,903 guinea pig) and 154,679 TG cells/nuclei across 5 studies (38,028 mouse, 116,651 human; Table S1). Neuronal and non-neuronal subtypes were then annotated and defined by their expression of known marker genes (Figure 1A-B, Table S2).

**Figure 1:**
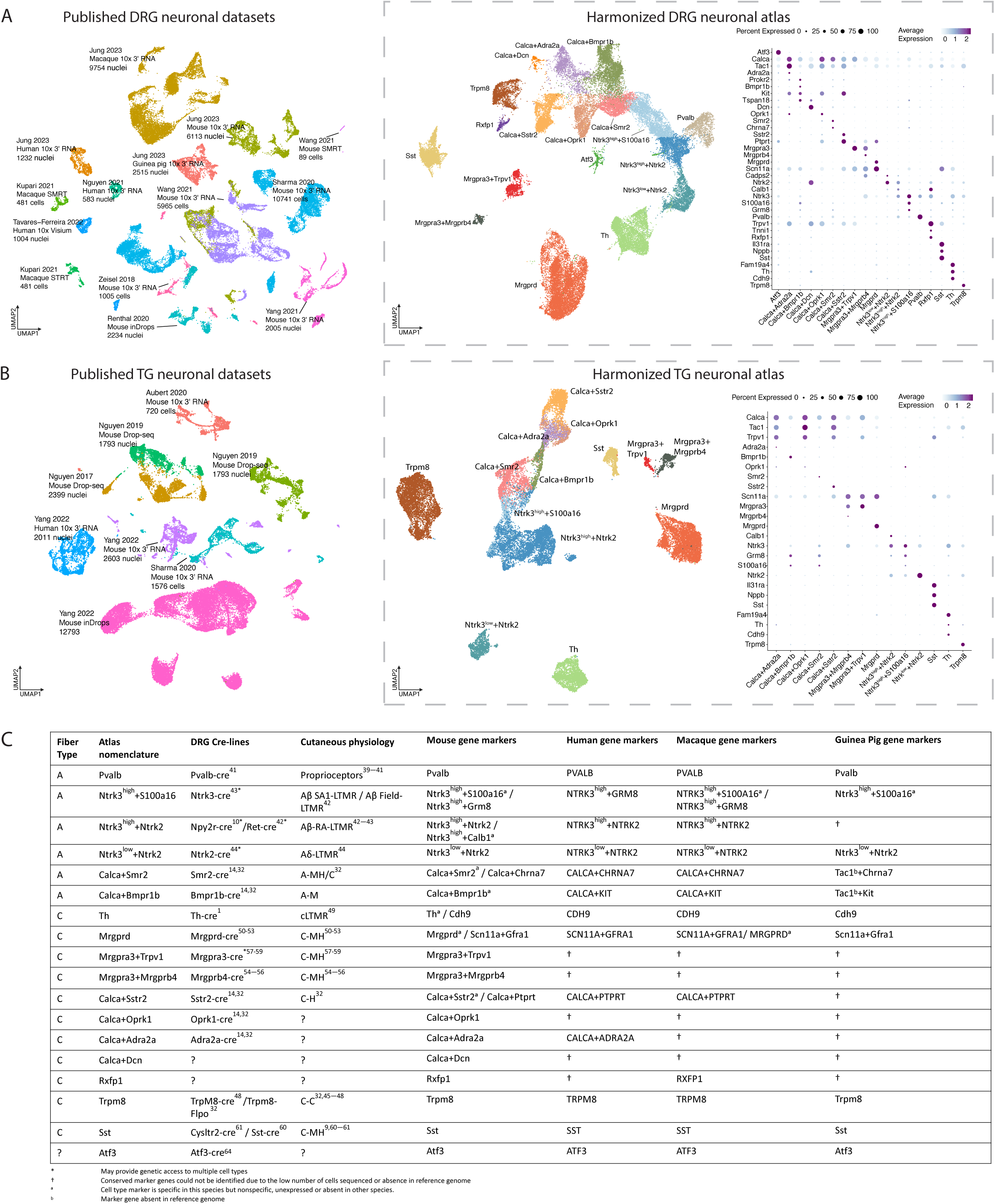
Integration of DRG or TG sc/snRNA-seq studies into harmonized neuronal atlases. A. *Integration of DRG neuronal sc/snRNA-seq datasets.* Left: Co-clustering of the 9 sc/snRNA-seq studies used in the harmonized neuronal DRG atlas. Each study’s citation, sequencing technology, species, and number of cells/nuclei sequenced are listed. Cells/nuclei are colored by study. Middle: UMAP projection of harmonized DRG neuronal atlas (44,173 cells/nuclei). Cells/nuclei are colored by their final cell type annotations in the harmonized atlas, which are named with their defining marker genes. Right: Dot plot of cell-type-specific marker gene expression. Dot size indicates the fraction of cells/nuclei expressing each gene and color indicates average log-normalized scaled expression of each gene. Abbreviations: SMRT = single molecule real time sequencing; STRT = Single-Cell Tagged Reverse Transcription sequencing. *B*. *Integration of TG neuronal sc/snRNA-seq datasets*. Left: Co-clustering of the 5 sc/snRNA-seq studies used in the harmonized neuronal TG atlas. Each study’s citation, sequencing technology, species, and number of cells/nuclei sequenced are listed. Cells/nuclei are colored by study. Right: UMAP projection of harmonized TG neuronal atlas (26,304 cells/nuclei). Cells/nuclei are colored by their final cell type annotations in the harmonized atlas. Right: Dot plot of cell-type-specific marker gene expression. Dot size indicates the fraction of cells/nuclei expressing each gene and color indicates average log-normalized scaled expression of each gene. *C*. *Somatosensory cell type nomenclature.* Atlas nomenclature along with fiber type, corresponding Cre-recombinase line, cutaneous physiology, and cell-type marker genes for each species. Abbreviations: LTMR = low threshold mechanoreceptor; C-M = C-fiber responsive to mechanical stimuli; C-H = C-fiber responsive to hot stimuli; C-C = C-fiber responsive to cold stimuli; C-MH = fiber responsive to mechanical and heat stimuli; A-M = A-fiber responsive to mechanical stimuli; A-MH/C = responsive to mechanical, heat and cold stimuli.

The harmonized DRG neuronal reference atlas contains 18 neuronal subtypes across 44,173 cells/nuclei and the harmonized TG neuronal reference atlas contains 14 neuronal subtypes across 26,304 cells/nuclei (Figures 1A-B). The annotations between the LIGER and Seurat pipelines agreed for 88% of the DRG neurons and 83% of the TG neurons, on average across cell types (Figures S1A; Table S3). Fifteen of the neuronal subtypes annotated in the harmonized atlases have at least one known function ascribed to them from rodent studies where direct genetic access to the cell type was obtained using knock-in mice expressing Cre recombinase (Figure 1C). These 15 neuronal subtypes include the Aβ-fiber subtypes *Pvalb* proprioceptors^39–,41^, Ntrk3^high^+Ntrk2 Aβ-rapid adapting (RA)-low threshold mechanoreceptors (LTMRs)^10,42^, Ntrk3^high^+S100a16 Aβ-field/slow adapting (SA)-LTMRs^43^, Ntrk3^low^+Ntrk2 Aδ-LTMRs^44^, Calca+Bmpr1b Aδ-high threshold mechanoreceptors (HTMRs)^14,32^, Calca+Smr2 Aδ-HTMRs^14,32^, and the C-fiber subtypes Calca+Sstr2 nociceptors^14,32^, Calca+Adra2a nociceptors^14,32^, Trpm8 cold thermoreceptors^45–48^, Th cLTMRs^49^, Mrgprd nociceptors^50–53^, Mrgpra3+Mrgprb4 cLTMRs^54–56^, Mrgpra3+Trpv1 itch-mediating neurons^57–59^, Sst itch-mediating neurons^59–61^, and an Atf3 cluster expressing injury-induced transcription factors such as *Atf3, Sox11* and *Jun*^26,62–64^. Eight Cre-lines that establish genetic access to these neuronal subtypes have been used to profile genetically labeled DRG neurons by bulk RNA-seq^16^; these transcriptomic profiles largely correlate (Pearson’s *r* > 0.5) with transcriptomic profiles of the respective neuronal subtype(s) in the harmonized DRG atlas (Figure S1B). We also observed three transcriptomically-defined clusters of C-fibers in the harmonized DRG atlas without known functions: Calca+Dcn, which expresses *Calca, Dcn,* and *Ntrk2,* Calca+Oprk1, which expresses *Calca, Oprk1,* and *Npy1r*, and Rxfp1, a rare subtype that is notable for expressing the highest levels of *Trpv1* in the DRG in addition to its marker gene, *Rxfp1* (Figure 1A).

With an exception of rarer neuronal subtypes (e.g. Mrgpra3+Mrgprb4, Mrgpra3+Trpv1, Calca+Oprk1, and Calca+Dcn), the DRG and TG neuronal subtypes defined in the harmonized atlases include cells/nuclei from all studies and species sampled (Figure S1D) that are transcriptomically similar (Jaccard indices > 0.6, Figure S1C). No human-specific clusters/neuronal subtypes were identified.

We observed the same neuronal subtypes in the harmonized TG atlas as the DRG atlas except for Atf3, Pvalb, Calca+Dcn, and Rxfp1, which we only observed in the DRG. The absence of detecting certain neuronal subtypes in the TG does not appear to be simply due to fewer or lower quality cells in the TG atlas as when we downsampled scRNA-seq data for the DRG, we still observe DRG-specific cell types (Figure S2A). Thus, it is possible that certain DRG cell types are much rarer or nonexistent in the TG, though additional TG scRNA-seq data are needed to confirm this observation. Nevertheless, cell-type-specific gene expression profiles (log_2_FC >0.5, adjusted p.value <0.05) of analogous TG and DRG neuronal subtypes are highly correlated (Pearson’s *r* >0.95, on average across cell types; Figure S2B) and ∼84% of TG neurons anchor with DRG neurons of the respective subtype (Figure S2C; see Methods).

While the harmonized DRG/TG neuronal atlases indicate that there are broadly similar neuronal subtypes between studies and species, there are notable nomenclature differences between individual studies that make it difficult to compare datasets. These nomenclature challenges are largely driven by technical factors such as the cell or nuclei dissociation protocol, number of cells/nuclei sequenced (Figure S2A), sequencing depth, or the single-cell technology used, all of which can affect the precise number of DRG or TG neuronal subtypes resolved in individual studies. Some groups label their sc/snRNA-seq clusters based on their associated cutaneous physiology; for example, *Calca+Smr2* cells have been described as Aδ-HTMRs^28^ or A-MH/C (A-fiber responsive to mechanical, heat, or cold stimuli)^32^. However, the physiology of transcriptomically-defined neuronal subtypes depends greatly on innervation target and not all cells sequenced innervate the skin. For example, Th neurons that innervate the skin function as cLTMRs and those which innervate adipose tissue modulate sympathetic activity^13^. For this reason, some groups describe their clusters of neurons simply by their expression of specific marker genes (e.g. *Nefh*-high clusters as NF1, or S100b/Ntrk3/Gfra1)^26,64^ or by a cluster number (e.g. C4)^18^. Here, we opted for a marker gene-based nomenclature because it is agnostic to function and scales in a consistent fashion as new subclusters emerge along with additional DRG/TG sc/snRNA-seq data. To assist in comparing similar transcriptomically-defined cell types across studies regardless of nomenclature, we designed the harmonized atlas metadata to facilitate rapid name conversion between the 15 DRG and 5 TG studies in both our data objects and our web resource (Figures S3A-D, https://harmonized.painseq.com).

### Harmonized mammalian somatosensory non-neuronal cell atlases

In the DRG, we identified 7 subtype-specific clusters across 187,387 cells/nuclei, and in the TG non-neuronal atlas, we identified 11 subtype-specific clusters across 88,155 cell/nuclei (Figure 2A – 2B; Table S1). The annotations between the LIGER and Seurat pipeline agreed for ∼90% of the DRG non-neurons and ∼93% of the TG non-neurons (Figure S1A). These include glial, endothelial, fibroblast and immune cells. With an exception of rarer cell types (e.g. immune cells), the harmonized non-neuronal cell types include cells/nuclei from all studies and species sampled (Figure S1C) that are transcriptomically similar (Jaccard indices > 0.6, Figure S1C). The major non-neuronal subtypes identified in the DRG are also present in the TG, though oligodendrocytes and astrocytes are uniquely present in the TG and may be derived from sequencing parts of the root entry zone where myelination switches form Schwann cells to oligodendrocytes. The TG is also more likely to contain meningeal fibroblasts than the DRG, though the fibroblast clusters in both the TG and DRG are likely a mixture of meningeal fibroblasts and those derived from epineurium, perineurium, and endoneurium.

**Figure 2:**
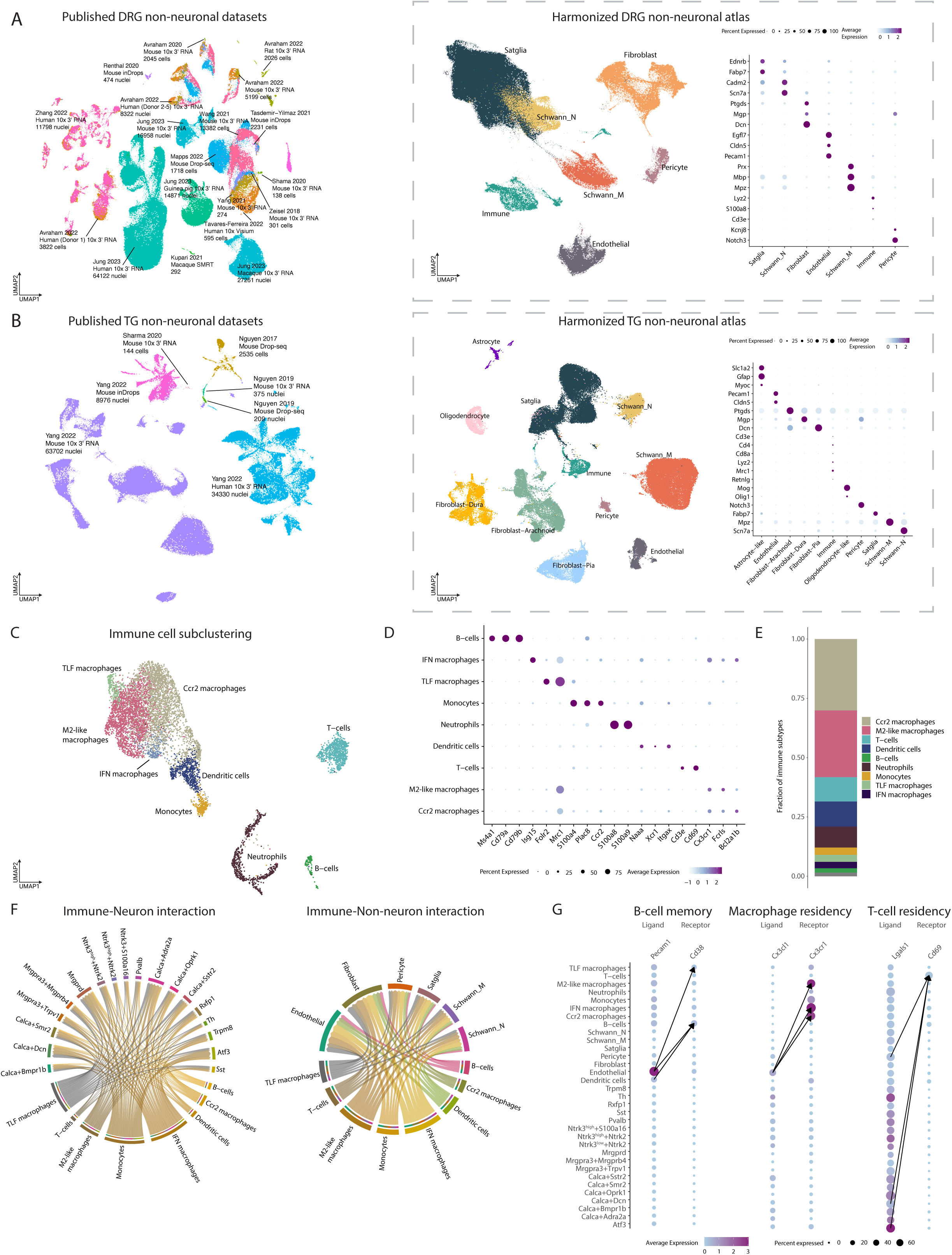
Integration of DRG or TG sc/snRNA-seq studies into harmonized non-neuronal atlases. A. *Integration of DRG non-neuronal sc/snRNA-seq datasets.* Left: Co-clustering of the 14 sc/snRNA-seq studies used in the harmonized neuronal DRG atlas. Each study’s citation, sequencing technology, species, and number of cells/nuclei sequenced are listed. Cells/nuclei are colored by study. Middle: UMAP projection of harmonized DRG non-neuronal atlas (187,383 cells/nuclei). Cells/nuclei are colored by their final cell type annotations in the harmonized atlas. Right: Dot plot of cell-type-specific marker gene expression. Dot size indicates the fraction of cells/nuclei expressing each gene and color indicates average log-normalized scaled expression of each gene. B. *Integration of TG non-neuronal sc/snRNA-seq datasets.* Left: Co-clustering of the 3 sc/snRNA-seq studies used in the harmonized non-neuronal TG atlas. Each study’s citation, sequencing technology, species, and number of cells/nuclei sequenced are listed. Cells/nuclei are colored by study. Middle: UMAP projection of harmonized TG non-neuronal atlas (88,155 cells/nuclei). Cells/nuclei are colored by their final cell type annotations in the harmonized atlas. Right: Dot plot of cell-type-specific marker gene expression. Dot size indicates the fraction of cells/nuclei expressing each gene and color indicates average log-normalized scaled expression of each gene. C. *Nine trascriptomically distinct immune cell types in DRG*. UMAP projection of DRG immune cells/nuclei (n = 8,178 cells/nuclei). D. *Marker genes used to annotate nine immune subtypes*. Dot plot of marker genes used to assign clusters to cell types. Dot size indicates the fraction of cells/nuclei expressing each gene and color indicates average log-normalized scaled expression of each gene. E. *Proportions of immune subtypes*. Fractions displayed represent the number of cells/nuclei for a given immune cell type out of the total number of immune cells. F. *Ligand-receptor interactions between immune cells and DRG neurons and non-neurons*. Predicted interactions bewteen ligands expressed by immune cells and their receptor pairs in DRG neurons (left) and non-neurons (right) (aggregated rank <0.005, see Methods). Color represents cell type. Arrow widths are proportional to the number of interactions between cell types. G. *Ligand-receptor interactions that may contribute to immune cell residency or memory.* Dot plot of ligand or receptor expression in DRG cell types. Dot size indicates the fraction of cells/nuclei expressing the ligand or receptor and color indicates average log-normalized gene expression. Arrows connect the cell-cell interactions that have the three highest ligand-receptor scores (see Methods).

We further classified glial cells/nuclei as satellite glia, myelinating Schwann cells, and non-myelinating Schwann cells. All clusters of satellite glia cells/nuclei express *Kcnj10* and *Glul*, while *Fabp7* and *Pou3f1* expression appears to be differentially expressed between the satellite glia clusters (Figure S4A), which is consistent with prior snRNA-seq studies of DRG satellite glia^21,65^. Myelinating Schwann cells (Schwann_M) express *Mpz*^66^, and non-myelinating Schwann cells (Schwann_N) express *Scn7a* and are located in the nerve (Figure S4B)^67^. Given the transcriptomic similarity between satellite glia and Schwann_N, spatial approaches that can differentiate between cells located in ganglia and nerve are likely better than sc/snRNA-seq for directly comparing these two glial cell types.

After the construction of the DRG non-neuronal atlas, we subclustered DRG immune cells and identified neutrophils, monocytes, B-cells, T-cells, dendritic cells, and macrophages based on established marker genes (Figure 2C-E, Table S2). While certain neutrophils and monocytes are predominantly found in circulation and dendritic cells are restricted to the tissue, the sequenced macrophages, B-cells, and T-cells could be a mixture of those derived from the circulation and DRG tissue. Indeed, resident macrophages have been well described in the DRG, and consistent with prior reports^68,69^, we observed in the harmonized DRG atlas that ∼34% of M2-like macrophages express the residency marker, *Cx3cr1*^69,70^, and ∼4% express the residency markers *Timd4/Lyve1/Folr2* (TLF macrophages; Dick *et al.*, 2019). Interferon (IFN) macrophages are inflammatory macrophages that express interferon responsive genes (*Isg15 and Irf7*) ^72–75^ and could be either DRG resident or circulating. While resident T-cells have been described in sensory ganglia during latent viral infections^76^, to our knowledge they have not yet been observed in naïve sensory ganglia. We thus asked whether residency markers associated T-cells in other tissues were also observed in DRG and found that *∼*48% of T-cells in the DRG atlas express residency markers (*Cd69* or *Itgae*)^77^. While tissue residency markers are less clear for B cells, we did observe that at least 57% of B cells are likely memory or mature B cells (*Cd27+* or *Cd20+*, respectively^78–80)^ and 22% are likely plasma cells (*Xbp1*+ or *Cd120+*^81,82^).

Taking advantage of the DRG immune cell atlas, we next performed ligand receptor interaction mapping across nine immune subtypes and 24 neuronal and non-neuronal cell types in the DRG (Figure 2F, Table S4) and asked which interactions might contribute to immune cell memory and residency within the DRG (Figure 2G; S4C). Memory B-cells express the receptor *Cd38* whose ligand (*Pecam1*) is predominantly secreted by endothelial cells. For resident macrophages, the interaction of *Cx3cl1* with macrophage receptor Cx3cr1 is an important driver of tissue residency^83,84^, and this appears to be largely mediated by the production of *Cx3Cl1* by DRG endothelial cells as well as several neuronal subtypes (Figure 2G). For resident T-cells, *Lgals1*, the classic ligand which drives T-cell residency by binding to Cd69, is equally expressed by nearly all neuronal and non-neuronal cell types. However, we noted pericyte-specific expression of *Myl9* (Figure S4C), a ligand whose binding to *Cd69* on T-cells has only recently been implicated in inflamed lungs ^85,86^.Our data suggests that *Myl9* expression by pericytes may be an additional activator of T-cell residency in sensory ganglia in addition to *Lgals1*.

In addition to exploring mechanisms of immune cell residency, we also noted ligand-receptor interactions between neuronal, immune and other non-neuronal DRG cell types that have been previously implicated in chronic pain, which could guide further mechanistic insight into these important neuro-glial-immune interactions (Figure S4D). Together, the harmonized DRG and TG cell atlases comprise a useful resource explore cell-type-specific gene expression profiles, neuro-glial-immune interactions, and to perform cross-study and cross-species comparisons of somatosensory cell types.

### Harmonized atlas has increased transcriptomic coverage compared to individual studies and improves cell type annotation of new human DRG snRNA-seq data

An important benefit of harmonizing previously published DRG and TG datasets is that additional cells can improve transcriptomic coverage within each cell type, which is particularly beneficial for rare cell populations in which there is currently insufficient coverage to detect lowly expressed genes with fidelity. Indeed, we observe that the harmonized DRG and TG neuronal atlases has greater (2.7-4.9 log fold change) transcriptomic coverage of lowly-expressed genes compared to individual studies alone (Figure 3A). This improved the detection of GPCRs, ion channels, transcription factors and peptides beyond any individual DRG and TG sc/snRNA-seq study (Figure 3B). The increased transcriptomic coverage was particularly helpful for identifying sensory cell types whose gene expression profiles may be affected by genomic variation associated with chronic pain in humans. Indeed, we found that genomic variation associated with chronic pain and headache occur in regions that may affect primate cell-type-specific expression profiles of Calca+Dcn, Calca+Mrgprb4, Ntrk3^low^+Ntrk2, Th, Calca+Bmpr1b and Neutrophil cell types (Figure S5A-B).

**Figure 3:**
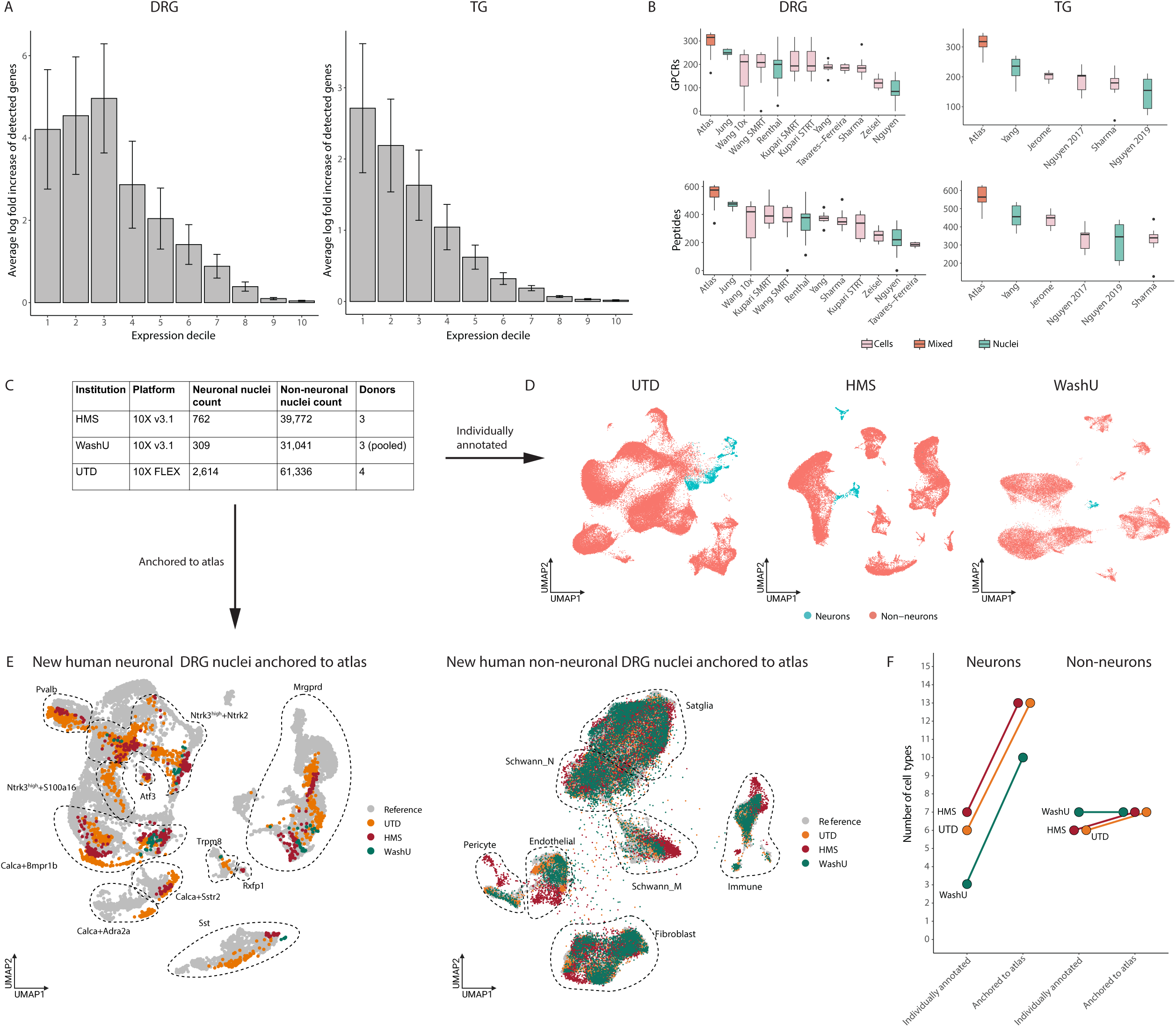
Harmonized reference atlas improves annotations of new human DRG snRNA-seq data. *A. Increased transcriptomic coverage of harmonized atlases compared to individual DRG and TG sc/snRNA-seq datasets.* Bar plots display the fold increase between the average number of detected genes per cell type in the harmonized atlas and the average number of detected genes in the respective cell type of each individual DRG or TG sc/snRNA-seq dataset. Expression deciles are defined using the atlas counts matrix where bin 1 represents genes with lowest expression and bin 10 represents genes with the highest expression. Error bars represent standard deviation. B. *Increased transcriptomic coverage of GPCRs and peptides in harmonized atlases compared to DRG and TG sc/snRNA-seq datasets*. Each box and whisker plot represents the distribution of the number of detected GPCRs or peptides per cell type in either the harmonized atlas or individual sc/snRNA-seq datasets. Studies with multiple species were combined. Dots represent outliers. C. *Number of nuclei sequenced in three new human DRG snRNA-seq datasets*. Human DRGs from 10 donors were sequenced at three different sites: University of Texas-Dallas (UTD), Harvard Medical School (HMS) and Washington University at St. Louis (WashU). D. *Neurons and non-neurons of the new human DRG snRNA-seq datasets.* UMAP visualization of 63,950 nuclei; WashU 31,350 nuclei; HMS 40,534 nuclei. Nuclei are colored by cell type. E. *Reference based cell type annotation of new human DRG snRNA-seq data.* UMAP projection of new human neuronal (left) and non-neuronal (right) data colored by institution. Cell types annotations of new snRNA-seq data after anchoring to the harmonized reference atlas are circled. Only nuclei with an anchoring score of >0.5 are displayed. Non-neuronal projection was downsampled to 100,000 nuclei to improve visualization. F. *Neuronal subtype resolution after anchoring new human DRG snRNA-seq data to reference atlas*. Plot of the number of DRG cell types that could be annotated individually compared to after anchoring to the reference atlas.

In addition to increased transcriptomic coverage and detection of rare cell populations, we also reasoned that the harmonized atlas could serve as a reference for annotating new DRG/TG sc/snRNA-seq datasets. This is particularly important in snRNA-seq studies of human sensory ganglia, as neurons are difficult to enrich prior to sequencing and are vastly outnumbered by non-neuronal cells. As such, individual datasets are often only able to sequence a small number of neurons, which can be difficult to annotate using traditional clustering approaches that are better suited for larger datasets. To directly test whether the harmonized atlas can be used as a reference to improve the annotation of new human DRG snRNA-seq data, we performed snRNA-seq of 135,834 nuclei from lumbar and thoracic DRGs from 10 donors at three different sites: Harvard Medical School (HMS; n = 40,534, 3 donors; Fig S6A), University of Texas-Dallas (UTD; n = 63,950, 4 donors; Fig S6B), and Washington University in St. Louis (WashU; n = 31,350, 3 donors; Fig S6C; Figure 3C). We analyzed and annotated these data using two parallel pipelines: 1) clustering/annotating each dataset individually (Figure 3D) or 2) individually anchoring each dataset to the harmonized DRG reference atlas (Figure 3E; see Methods). Despite the inherent limitations of cell type annotations in the absence of an established ground truth, ∼87% of the cell type annotations made using the reference atlas agreed with the broader cell type assignment made by clustering the data separately (Figure S6E, see Methods). Moreover, the reference atlas was able to resolve and annotate ∼2-3 times as many DRG neuronal subtypes than when clustering/annotating each dataset individually (Figure 3F; Table S1; Figure S6E). Our findings suggest that the reference atlas can not only simplify cell type annotations of new human snRNA-seq datasets but also improve the granularity of these annotations, particularly for rarer cell types that cannot be resolved when clustering smaller datasets individually.

### Comparison of mouse and human sensory ganglia

Harmonizing somatosensory sc/snRNA-seq datasets from multiple species enables direct cell-type-specific transcriptomic comparisons across species, which is particularly important for understanding similarities between human and mouse sensory ganglia, as mice are often used to interrogate cell type function and model pathology (e.g. neuropathic pain).

We thus compared the cell-type-specific gene expression across species (log_2_FC>0.5, adj. p-value < 0.05 relative to all other TG/DRG neuronal subtypes in the TG/DRG). We observed statistically significant overlap between the neuronal subtype-specific genes expressed in mouse DRG and those expressed in human DRG across all neuronal subtypes (P < 0.01, hypergeometric test; Figure 4A-B; Table S5). We also observed significant overlap between neuronal subtype-specific gene expression in mouse and human TG as well as between non-neuronal subtype-specific gene expression in mouse and human DRG/TG. Some of these evolutionarily conserved neuronal and non-neuronal cell-type-specific genes include well known neuropeptides (e.g. *ADCYAP1* in Calca/Sstr2 neurons), ion channels (e.g. *TRPM8* in Trpm8 neurons), GPCRs (e.g. *EDNRB* in Satellite glia neurons), and transcription factors (*ATF3* in Atf3 neurons; Table S6). The evolutionarily conserved expression patterns of these genes suggest that functional studies of these genes in mice DRG/TG could be predictive in many cases to their function in humans.

**Figure 4:**
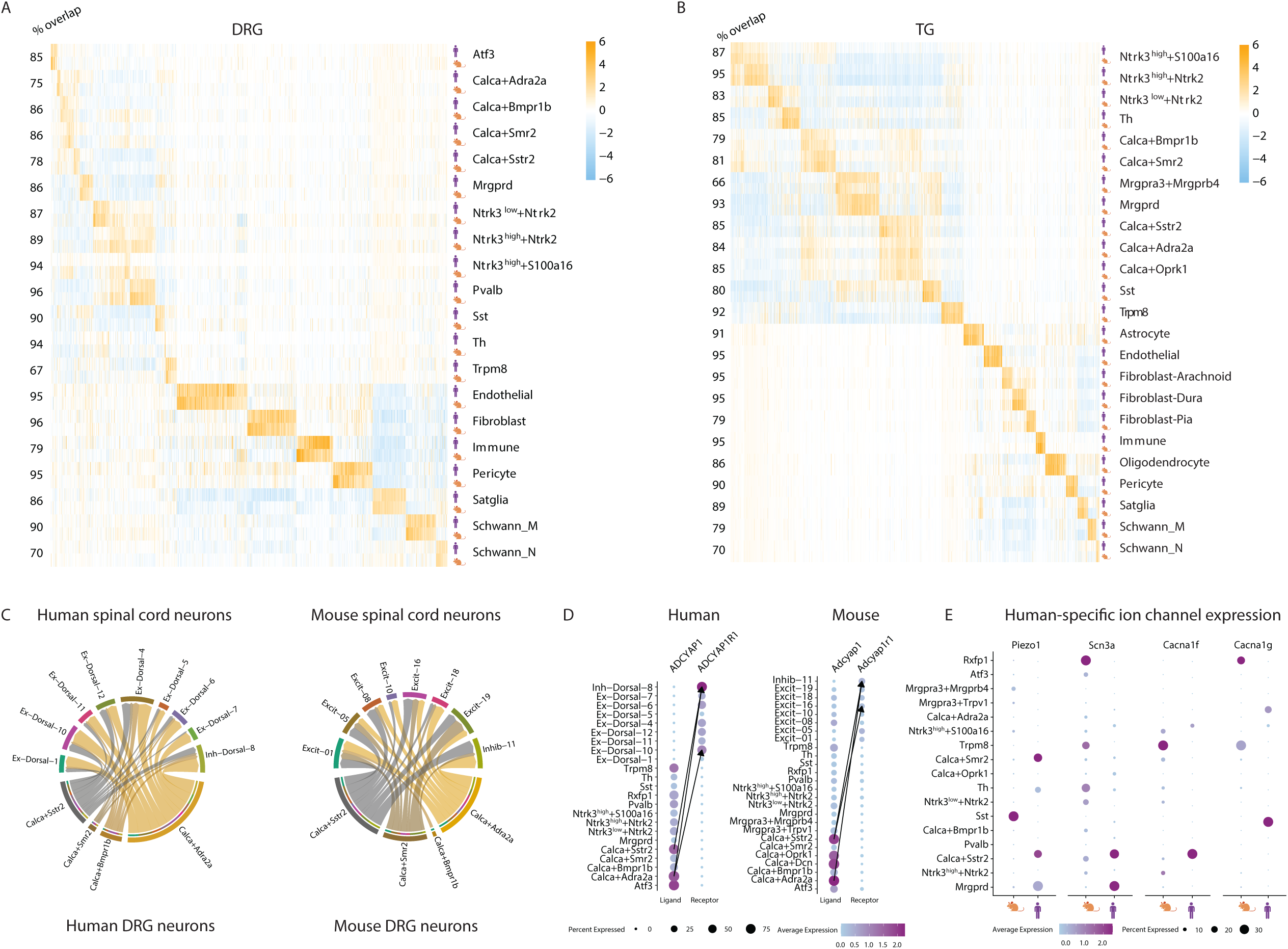
Cell type-specific gene expression patterns among human and mice. *A-B. Overlap of cell-type-specific gene expression between mouse and human cell types.* Displayed are the log_2_FC values of cell-type-specific genes (columns) of either human or mouse (log_2_FC>0.5, adjusted p.value < 0.05) when comparing expression in one cell type compared to all other DRG cell types of the same species. Numbers to the left of rows (% overlap) represent the percent of human cell-type-specific genes that overlap with mouse cell-type-specific genes. Cell types with fewer than 20 human nuclei were not displayed. *C. Ligand-receptor interactions between DRG Calca+ neurons and spinal cord dorsal horn neurons.* Predicted interactions between ligands expressed by human and mouse DRG neurons and their receptor expressed by dorsal horn neurons (aggregated rank <0.005, see Methods). Arrow widths are proportional to the number of interactions between cell types. *D. Similar cell types mediate the Adcyap1 and Adcyap1r1 interaction in human and mouse.* Dot plot of ligand expression in human and mouse DRG neurons or receptor expression in spinal cord neurons. Dot size indicates the fraction of cells/nuclei in each cell type expressing the ligand or receptor and color indicates average log-normalized gene expression. Arrows connect the cell-cell interactions with the three highest ligand-receptor scores (see Methods). *E. Species-specific expression of ion channels.* Dot plot displays expression of *Piezo1*, *Scn3a*, *Cacna1f*, *Cacna1g* in human and mouse DRG neuronal subtypes. Dot size indicates the fraction of cells/nuclei expressing each gene and color indicates average log-normalized expression.

Somatic pain transduction requires signaling from DRG neurons to the dorsal horn of the spinal cord, so we next compared human and mouse ligand receptor interactions between DRG neurons in the atlas to recently published snRNA-seq spinal cord data^38,87^. We noted that in both mouse and human, Calca+Adra2a and Calca+Sstr2 neurons express the largest number of ligands that have receptor pairs with dorsal horn neurons, and that these interactions occur with similar dorsal horn neuronal populations in both species (Inhib-11/Inh-Dorsal-8, Excit-01/Ex-Dorsal-1, Ex-Dorsal-4/Excit-05, Excit-16/Ex-Dorsal-10; Figure 4C; Table S7 for all LR pairs) with known roles in mechanical allodynia^88^ and sensorimotor reflex to pain^89^. As an example, we observed that in both mouse and human, Calca+Adra2a and Calca+Sstr2 neurons express *Adcyap1* and Inhib-11/Inh-Dorsal-08 express its receptor *Adcyap1r1* (Table S7; Figure 4D).

While our cross-species analysis of the harmonized DRG/TG atlases support broad transcriptomic similarity between human and mouse, there are also notable differences between their cell-type-specific transcriptomes. Indeed, we observed >500 cell-type-specific genes in the DRG or TG that are expressed significantly more in human than in mice (log2FC>0.5, adj. p-value<0.05 human vs. all non-human species; Table S8). For example, consistent with prior reports^25,28,29^, several neuropeptides that are restricted to specific cell types in mice are expressed more broadly across neuronal subtypes in both the DRG and TG in humans. The increased transcriptomic coverage of the harmonized atlas also allowed us to explore species differences in more lowly expressed genes. We observed that *Piezo1*, which is predominantly expressed by SST neurons and contributes to alloknesis in mice^59^, is lowly expressed across multiple DRG neuronal subtypes in human. Conversely, we observed several ion channels that are expressed broadly in mice but are highly restricted to nociceptive subtypes in human (e.g. *Cacna1f*), and which may warrant additional study in human neurons. Taken together, our analyses indicate that despite broad similarity between mice and humans with regard to expression of cell type identity genes, notable species differences exist in the expression of functionally important genes that may help specialized sensory neurons interact with specific environmental niches.

### Evolutionary insights from sensory ganglia transcriptomes across non-mammalian animal models

We reasoned that the harmonized DRG atlas could also be used as a reference to compare gene expression in sensory ganglia cell types across a wider evolutionarily landscape of animal models. While we were able to obtain published sc/snRNA-seq data from mammalian vertebrates (e.g. mouse^17,18,20,23,27,31,64^, rat^22^, guinea pig^31^, and macaque^24,31^ DRGs) and invertebrates (*Drosophila melanogaster* chordotonal organ^90^ (Figure S8A) and *C. elegans*^91^), we noted that non-mammalian vertebrate sensory ganglia have not been previously characterized at single-cell resolution.

As axolotls are both non-mammalian vertebrates and commonly used models to study nerve and limb regeneration^92,93^, we sequenced 4,848 DRG cells (12 DRGs total from 2 animals; Figure 5A). We annotated 1,817 cells as neurons (Figure 5B-E) and 3,031 as non-neurons (Figure 5F-G; see Methods). The neurons were separately subclustered (Figure 5B). As with mammals, we observed an NEFH high and NEFH low population (Figure 5C), suggesting the presence of large to medium diameter myelinated A-fibers and small diameter unmyelinated C-fibers. The putative A-fiber population consisted of distinct clusters that anchored to mammalian Pvalb and Ntrk3^high^+Ntrk2 A-fiber populations (Figure 5D; see Methods). The putative C-fiber neurons separated into two peptidergic neuron populations, C-X1 and C-X2 that express *Tac1, Adcyap1, Adra2a, Trpv1 and Trpm8*, and display the transcriptomic similarity to mammalian Calca+Adra2a, Calca+Bmpr1b, Rxfp1 and Trpm8 neurons (Figure 5D). A notable difference between axolotl and mammamlian C-fibers is that *Trpm8* is expressed in >40% of C-fibers in axolotols compared to ∼3% in mammals (Figure 5E). Additionally, ∼17% of axolotl C-fibers co-express *Trpm8* and *Trpv1*, which is higher than the ∼0.25% of mammalian C-fibers that co-express *Trpm8* and *Trpv1.* Together, these data suggest an expansion of C-fibers with the ability to detect both cold and heat in the axolotl, which may provide these cold-blooded amphibians a more refined ability to perceive temperature^94,95^.

**Figure 5:**
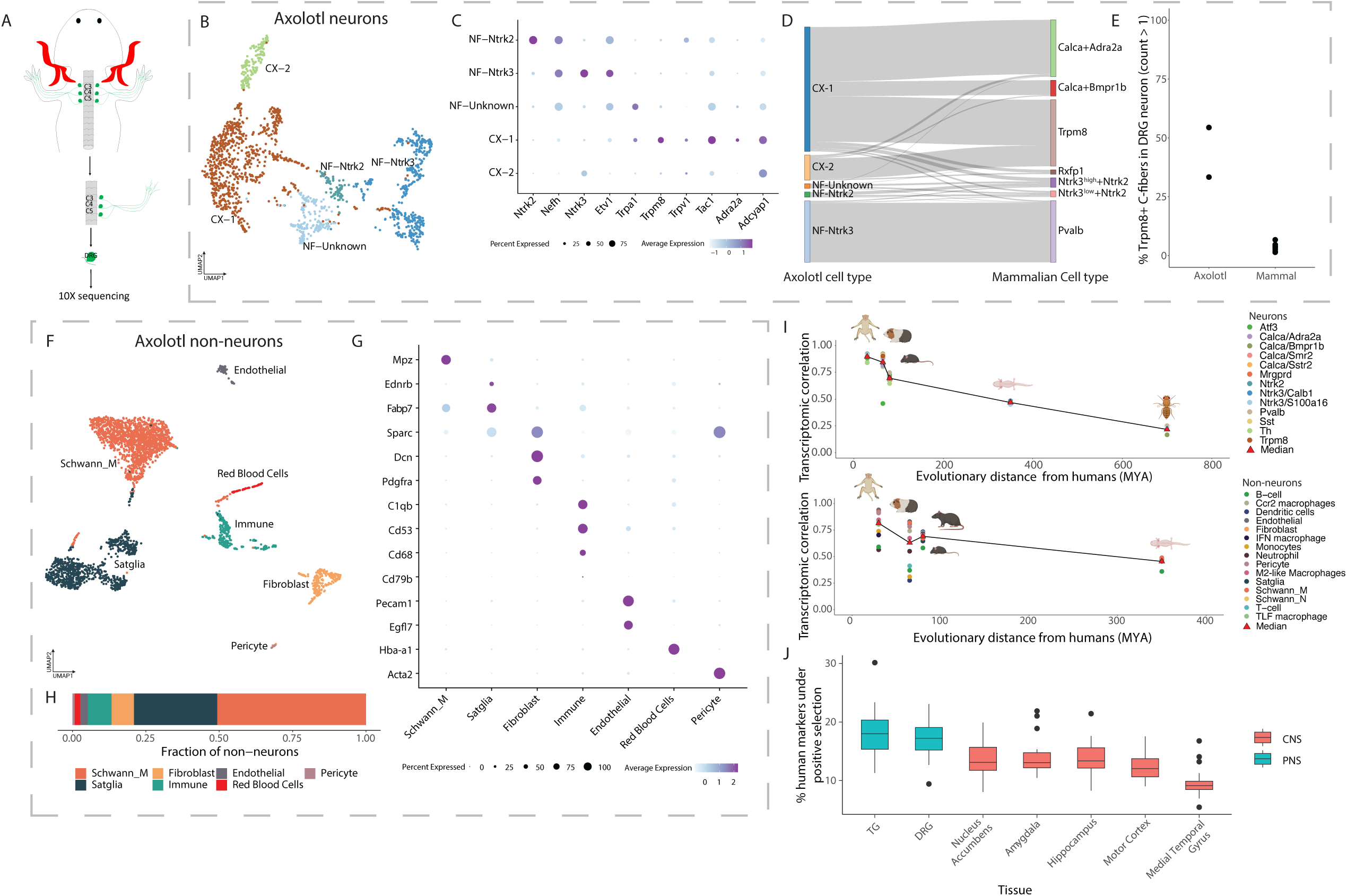
Transcriptomic similarity between axolotl and mammalian DRG cell types. A. *Axolotl DRG collection schema for scRNA-seq.* Cervical DRGs (C3, C4 and C5) were collected (2 animals; total of 6 DRGs/animal), freshly dissociated and transcriptomically profiled by scRNA-seq. B. *Axolotl DRG neuronal subtypes*: UMAP plot of 1,817 neurons from the axolotl DRG that form 5 transcriptomically distinct subtypes. A-fiber subtypes are denoted using “NF” and C-fiber subtypes are denoted using “CX”. Cells are colored by cell type. C. *Expression of marker genes used to annotate axolotl neuronal cell types.* Dot plot of marker gene expression in axolotl DRG neurons. Dot size indicates the fraction of cells expressing each gene and color indicates average log-normalized expression. D. *Transcriptional similarity between axolotl and mammalian DRG neurons.* Sankey plot displays the annotation of each axolotl DRG cell after anchoring to the DRG reference atlas. Only cells with anchoring scores greater than 0.5 are diplayed. E. *More axolotl C-fibers express Trpm8 compared to mammalian C-fibers.* Fractions display the number of C-fibers that express *Trpm8* over the total number of C-fibers in axolotols or the DRG neuronal reference atlas. F. *Axolotl DRG non-neuronal subtypes.* UMAP plot of 3,031 non-neuronal cells from the axolotl DRG that form 7 transcriptomically distinct subtypes. Cells are colored by cell type. G. *Expression of marker genes used to annotate axolotl non-neuronal cell types.* Dot plot of marker gene expression in axolotl DRG non-neurons. Dot size indicates the fraction of cells expressing each gene and color indicates average log-normalized expression. H. *Cell type proportions of axolotol non-neurons.* Proportions displayed are a ratio of the number of axolotl non-neuronal DRG cells for a given cell type to the total number of non-neuronal cells. I. *Transcriptomic correlation with DRG cell types decreases over evolutionary distance.* Plot displays for each species, the correlation between the average expression (log normalized counts) of each gene in each cell type to the average expression (log normalized counts) of each gene in the corresponding human cell type (y-axis), plotted against the evolutionary distance from humans (millions of years ago [MYA]; x-axis). Red triangles represent the median correlation for each species across all cell types. For display purposes, cynomolugus macaque and rhesus macaque were grouped together as well as mouse and rat. J. *Positive selection of cell-type-specific gene expression in PNS and CNS.* Boxplots represent the number of human marker genes (log_2_FC>0.5, adjusted p.value < 0.05 relative to other nuclei in the same species) per cell type that are evolving under positive selection (see Methods). PNS = peripheral nervous system; CNS = Central nervous system.

We next compared cell-type-specific transcriptomic profiles of sensory ganglia from human, non-human primates, mouse, rats, guinea pig, axolotls, and *Drosophila* by calculating the correlation between the average normalized counts for each corresponding cell type between each species. Consistent with a prior report^31^, we found that as evolutionary distance from humans increased, the transcriptomic correlation with the most similar non-human sensory cell type decreased (Figure 5I).

Genes expressed by cells at the interface between organism and environment (e.g. immune cells) are more likely to evolve under positive selection (the process by which advantageous variants are selected for during evolution) than those not in direct contact with the environment^96–99^. We therefore asked whether DRG/TG cell-type-specific genes are also more likely to be under positive selection than other central nervous system cell types. Indeed, we observed that human DRG/TG cell-type-specific genes are approximately twice as likely to be under positive selection than those expressed by a range of CNS cell types (Figure 5J; Table S9). The set of cell-type-specific genes evolving under positive selection includes genes with previously established somatosensory functions such as ion channels (*TRPM8* in Trpm8 neurons^97^), neuropeptides (*CALCB* in Calca+Smr2 neurons), and GPCRs (*NPY2R* in Sst neurons). Future research is required to understand how specific environmental niches may drive DRG/TG cell-type-specific genes to evolve under positive selection.

The harmonized DRG/TG atlases along with the new multi-species scRNA-seq data and analyses presented here not only provide a valuable resource for studying cell-type-specific gene expression in sensory ganglia across studies and species, but also provide novel insight into species differences in somatosensory cell identity and the evolutionary mechanisms that contribute to this process.

## Discussion

By harmonizing 19 sc/snRNA-seq studies from sensory ganglia of human, mouse, rat, guinea pig and macaque, we constructed a DRG/TG reference atlas that enables comparisons across studies and species with improved transcriptomic coverage and cell type resolution than any study offers individually. We have also built a web portal (https://harmonized.painseq.com) to facilitate access to this resource across the research community.

The exponential increase in sc/snRNA-seq datasets from sensory ganglia over the last few years has yielded incredible new molecular insights^11,17–24,24–26,26–32^ but has also led to nomenclature challenges between studies as new clusters/subtypes emerge with additional sequencing data from a range of studies, conditions, and species^11,100^. A key application of the atlas presented here is to harmonize cell type nomenclature across studies; indeed, both the metadata and webtool enable rapid conversion of cell type names between studies as well provide a reference to which new sc/snRNA-seq studies can be rapidly anchored and annotated.

We opted to base the reference atlas nomenclature on DRG marker genes because it allows for a consistent naming strategy as new clusters/subtypes/marker genes emerge with future transcriptomic, epigenomic, and spatial data. Moreover, a marker gene-based nomenclature remains agnostic to cell type function, which is known to vary widely depending on projection target^13,14,26,32^. We opted to use mouse DRG marker genes for the cell type nomenclature because mouse is the most commonly used mammalian model system, a majority of the current atlas is comprised of cells/nuclei from mice, and Cre lines driven by these marker genes are available for most subtypes to help characterize their function in a range of tissues^14,32,101^. However, we concede this nomenclature strategy is imperfect. In addition to failing to incorporate historical nomenclatures derived from studies of cutaneous physiology, certain mouse marker genes are not present in other species (Table S10). For example, the Calca+Smr2 subtype is present across mammals, but humans, along with most mammals, have no known 1:1 ortholog for the mouse *Smr2* gene. Thus, we envision that as new sc/snRNA-seq data are generated from human^35^ and other species, this atlas and its nomenclature will need to be updated to reflect the emergence of new cell types until all sensory ganglia cell types have been sequenced to a point of saturation. In the interim, we feel that the harmonized atlas and proposed nomenclature provide a straightforward way to communicate about the same transcriptomic cell types across studies regardless of the names each study uses to describe their data.

In total, we observed 29 transcriptomically-defined cell types in the cross-species harmonized DRG/TG cell atlas, 18 neurons and 11 non-neurons. Consistent with prior reports from the cerebral cortex^102,103^, scRNA-seq data was superior to snRNA-seq data for resolving neuronal subtypes. While there is sufficient information from nuclear mRNA to resolve all subtype identities by anchoring to the reference atlas, we believe that generating new scRNA-seq over snRNA-seq data across a range of species will maximize the ability of future reference atlases to resolve rarer subtypes.

We observed evidence of broad transcriptomic conservation in sensory ganglia cell types across species ranging from *Drosophila melanogaster* to humans, though the correlation between cell-type-specific gene expression profiles decreased with evolutionary distance form human. For example, similar classes of transcriptomically-defined DRG cell types are present in axolotls and mammals (Figure 5E), but there also notable differences including the expansion of C-fiber subtypes that co-express channels responding to heat (*Trpv1*) and cold (*Trpm8;* Figure S8C). These channels are rarely expressed in the same neurons in mammals.

None of the neuronal subtypes we identified in the harmonized atlas or our new human DRG snRNA-seq data were unique to human, which indicates broad conservation of cell-type-specific gene expression profiles across mammalian species (Figure 4A-B). This finding is consistent with previous snRNA-seq study of human and mouse TG^29^, and sc/snRNA-seq of human, macaque, and mouse DRG^24,31^. However, deeper sequencing of human DRG will provide improved resolution of human sensory ganglia and may reveal additional subtypes^35^. Indeed, we observed that human cell-type-specific gene expression in DRG/TG is enriched for genes undergoing positive selection compared to CNS subtypes (Figure 5J), which highlights that human-specific features likely warrant further investigation. Fortunately, significant advances in culturing peripheral human DRG neurons^104^ and engineering human iPS nociceptors^105–107^ together should allow for mechanistic exploration of human-specific features directly in human cells.

With our increased coverage of physiologically relevant genes (Figure 3B), we found lowly expressed and/or understudied genes that may be attractive candidates for new pain therapeutic targets because they are expressed in the same nociceptor subtypes in both humans and mice. One example is the GPCR *Gpr26,* which is an orphan G_s_-coupled GPCR that has sequence similarity to the purinergic GPCR P2YR^108^, and is expressed in the Trpm8 cell type in both mouse and human (Figure S7A-B). Additionally, with our increased cell type resolution, we were able to characterize nine immune cell types including two tissue resident immune cell types for both humans and mice (Figure 2C-G), warranting future work in understanding how DRG/TG injury and infection can lead to immune residency in glial structure. The increased transcriptomic coverage was particularly helpful for identifying sensory cell types whose gene expression profiles may be affected by genomic variation associated with chronic pain in humans. Indeed, we found that genomic variation associated with chronic pain and headache occur in regions that are likely to affect cell-type-specific expression profiles of Calca+Dcn, Mrgpra3+Mrgprb4, Ntrk3^low^+Ntrk2, Th, Calca+Bmpr1b and Neutrophil cell types (Figure S5A-B). Our findings are broadly consistent with a previous report^24^ that *Calca*+ and *Mrgpra3*+ cells contribute to chronic pain heritability, though technical differences in our analytical approaches make it difficult to directly compare our study with Kupari et al., 2021.

The harmonized DRG/TG atlases and new multi-species data presented here enable comparisons across studies and species with improved transcriptomic coverage and cell type resolution. However, this resource should be considered in the context of its limitations. There are multiple strategies for harmonizing sc/snRNA-seq datasets that can affect cell type annotation^33,34,109^. We opted for an approach in which cells/nuclei are excluded from the atlas if they are inconsistently annotated (∼7-17% of cells/nuclei) across two independent and benchmarked integration pipelines^33,34^. While incorporating additional integration strategies could further strengthen cell type assignment confidence of existing data, we believe that new data, especially neuronal scRNA-seq data from non-rodent species, will provide the best opportunity to confidently identify and profile the complete transcriptomes of rare sensory ganglia cell types. Future profiling of sensory ganglia cell types should also include epigenomic, spatial, projection target, and physiology across a range of conditions and species, all of which will contribute to the definition of ‘cell types’^110^ in future versions of this atlas.

## Supporting information

Table S1 - Dataset summary

Table S2 - Marker genes

Table S3 - Seurat-LIGER Comparison

Table S4 - Ligand-receptor all cell types

Table S5 - Overlapping markers

Table S6 - Human Ligand Receptor

Table S7 - Mouse Ligand Receptor

Table S8 - Genes under positive selection

Table S9 - Nomenclature across species

Table S10 - Human donors

Table S11 - Axolotl HCR probes

## Acknowledgements

This work was primarily supported by the Migraine Research Foundation (W.R.), Burroughs Wellcome Fund (W.R.), Barry Family Harvard Stem Cell Institute Award (J.L.W), and the National Institute of Neurological Disorders and Stroke (W.R & C.J.W. U19NS130617, R01NS119476 (W.R.), T. D. U19NS130608 (T.J.P), U19NS130607 (R.W.G)). W.R. also receives support from the National Institute of Neurological Disorders and Stroke (K08NS101064 and), National Institute of Drug Abuse (DP1DA054343), National Eye Institute (U01EY034709), Teva Pharmaceuticals, and BWH Women’s Brain Initiative Neurotechnology studio. The authors would like to thank donors and their families, without whom much of this study would not be possible. We would also like to thank members of the Renthal, Woolf, Price, Gereau and Whited labs as well as Kaylee Gentry, Jordan Sicherman, Areesha Salman, Shigeru Saito, Shreejoy Tripathy, and Jesse Gillis for their helpful feedback throughout the study.

## Data and Code Availability

Processed data are available at harmonized.painseq.com. Raw and processed data will be available within the Gene Expression Omnibus (GEO) repository (www.ncbi.nlm.nih.gov/geo). Custom R scripts will be available on GitHub at github.com/Renthal-Lab/harmonized_atlas.

## Materials and Methods

### Publicly available scRNA-seq somatosensory data collection

Original datasets were downloaded from GEO (Table S1). If original cell type annotations were not provided, we followed the annotation protocol described by the original study. Any unresolved clusters would be labelled as unknown. If the data provided on GEO was not quality controlled for number of features and counts or for proportion of mitochondrial counts, we applied quality control filters based on the sequencing platform and protocol. For the purposes of integration, human, guinea pig, and macaque were converted to mouse gene names using Ensembl’s biomart v109^111^.

### Integration of publicly available scRNA-seq somatosensory data

To alleviate study-specific batch effect in integrating and clustering the sc/snRNA-seq data, we used two different computational tools for data integration: Seurat v4 and LIGER^36,37^. While both tools overlap in using Louvain clustering and UMAP dimensionality reduction, they differ in how study-specific variable genes are selected, how the consensus of variable genes across all studies are selected and the neighborhood graph construction. If we labelled a cell with the same cell type with both LIGER and Seurat, then that was more likely to be the correct cell type for the cell. Similarly, if we annotated a cell differently when using LIGER than in Seurat, we removed the cell from the final atlas. Final UMAP projection presented in this paper are Seurat-based unless otherwise stated.

Unless otherwise stated, all Seurat integration and clustering used 2000 variable genes, and 30 PCs and all LIGER integration used 2,000 variable genes and 30 k factors for non-negative matrix factorization. All integration analyses integrated across “datasets” which we defined as a specific study, species, and platform because a single study can have data from different platforms and species. Integration anchors were chosen using scRNA-seq datasets. Minimum number of features was 1000 for DRG neuronal, DRG non-neuronal and TG non-neuronal integration, while minimum number of features for TG neuronal integration was 400. Maximum mitochondrial fraction was 10%.

After the initial round of clustering, we separated neuronal and non-neuronal clusters based on Rbfox3 and Sparc expression (log_2_FC>0.5 and adjusted p.value <0.05 relative to all other clusters in the same UMAP space; Table S2). We then reintegrated and reclustered neurons and non-neurons, then neurons and non-neurons were annotated based on marker gene expression (Table S2). If a cluster contained cells/nuclei that was 90% from a single dataset, we considered this to be a study-specific batch effect and we removed it from our final analysis. Clusters were labelled doublets based on overlapping neuronal and non-neuronal marker gene expression (Table S2). With the exception of Seurat and LIGER annotations for TG neurons, we then subclustered Nefh^high^ cells/nuclei and Nefh^low^/Calca cells/nuclei from the harmonized atlas to annotate with A-fiber markers or peptidergic C-fiber markers.

For Seurat annotations of TG subclustered Nefh^high^ cells/nuclei and Nefh^low^ /Calca, we used Seurat’s label transfer feature (“anchoring”) to cluster Nefh^high^ cells/nuclei and Nefh-low/Calca cells/nuclei from the harmonized atlas to the Nefh^high^ cells and Nefh^low^/Calca cells from Sharma and colleagues’ study ^23^. Briefly, FindTransferAnchors(reduction= “cca”) was used to identify anchors between identify anchors between Nefh^high^ cells/nuclei and Nefh^low^/Calca cells/nuclei from the harmonized atlas to the Nefh^high^ cells and Nefh^low^/Calca cells from Sharma and colleagues’ study. TransferData() was used to transfer Sharma and colleagues’ labels to the Nefh^high^ cells/nuclei and Nefh^low^/Calca cells/nuclei. Variable genes were identified from the merged dataset, and PCA and UMAP were run to generate new UMAP coordinates.

Our protocol to integrate, cluster and annotate TG neurons using LIGER was similar to our Seurat-based protocol with a key difference in how we annotated Nefh^high^cells/nuclei and Nefh^low^/Calca cells/nuclei. These cells/nuclei were separated from other neurons, reintegrated and reclustered using LIGER. Final cluster assignments were given based on which cell type from Sharma and colleagues made up the largest proportion of that cluster.

All code used to integrate data will be made available on GIT after manuscript acceptance. Final harmonized atlas objects will be made available after manuscript acceptance. Final Seurat object are made available for exploration on an online Shiny app constructed using ShinyCell^112^.

### Immune cell subclustering

DRG immune cells were separated, reintegrated and subclustered after construction of the DRG non-neuronal atlas. We then reintegrated and reclustered neurons and non-neurons, then neurons and non-neurons were annotated based on marker gene expression (log_2_FC>0.5 and adjusted p.value <0.05 relative to all other clusters in the same UMAP space; Table S2).

### Identifying shared and species-specific expression in the somatosensory atlas

The cells/nuclei for each species were separated for each of the final atlases (DRG neurons and non-neurons, and TG neurons and non-neurons), and then FindAllMarkers was run across all final subtypes. For each subtype, we identified whether a gene was differentially expressed (log_2_FC>0.5 and adjusted p.value <0.05 relative to all other subtypes in the same species). Cell types with less than 20 human nuclei were excluded. DRG mouse data was downsampled to have the ratio between the DRG mouse cells and DRG human match the ratio between the TG mouse data and the TG human data (Figure 4A).

### Ligand receptor pair analysis

Ligand-receptor pair analysis was performed using LIANA ^113^. The DRG non-neuronal and neuronal atlases were combined for Figures 2F-G and S4C. We then ran LIANA using default settings using their mouse or human consensus database and we filtered ligand-receptor interactions with an aggregated rank <0.005 (Figure 2F). The SingleCellSignalR scores generated by LIANA was used for visualization in Figure 2G and S4C. Predicted mouse and human interactions were combined for visualization.

For our ligand-receptor pair analysis between human/mouse DRG neurons and human/mouse spinal cord neurons, we used spinal cord datasets published by the Levine lab ^38,87^. Human and mouse spinal cord cell types were subsetted for the cell types found in both species as described in the Yadav et al., 2023 study. We then merged the human spinal cord neurons with the human DRG neurons, and merged the mouse spinal cord neurons with the mouse DRG neurons. We then ran LIANA on the merged objects using default settings using their human or mouse consensus database and we filtered ligand-receptor interactions with an aggregated rank <0.005. The SingleCellSignalR scores generated by LIANA was used for visualization in Figure 4.

### Cell type Jaccard score

We calculated Jaccard scores using cell types that are clearly defined clusters across both single cell and single nuclei RNA-seq datasets. For neurons, these cell types were Sst, Mrgprd, Pvalb, Th and Trpm8. For non-neurons, these cell types were Schwann_M, Fibroblast, Endothelial and Pericytes.

For each study with a reported Sst cell type, we calculated a Jaccard similarity index where Set A is defined as all cells/nuclei labeled as Sst from a given study in the harmonized atlas and Set B is defined as all cells/nuclei labeled as Sst in the harmonized atlas. The Jaccard index was calculated as the intersection of Set A and B divided by the union of Set A and B. We repeated the same measurements for neuronal atlases with reported Mrgprd, Pvalb, Th and Trpm8 cell types and for the non-neuronal atlases with reported Schwann_M, Fibroblast, Endothelial and Pericyte cells.

### Physiologically relevant genes

Mouse gene names for GPCRs were taken from the IUPHAR/BPS guide to PHARMACOLOGY database^114^. Mouse gene names for peptides were taken from CellTalkDB^115^. Gene names were intersected with detected genes (average count > 0) for each cell type.

### TG and DRG label transfer

To directly compare the TG atlas to the DRG atlas, we used Seurat v4 to “anchor” the TG atlas to the DRG atlas. FindTransferAnchors(reduction = ‘‘cca’’) in Seurat was then used to identify anchors between DRG and TG data. TransferData() was used to transfer DRG subtype labels to each nucleus/cell in the TG atlas. TG nuclei with anchoring prediction score < 0.5 were excluded from the dataset. Variable genes were identified from the merged dataset, and PCA and UMAP were run to generate new UMAP coordinates.

### Mice

For in situ experiments, adult (8-12 week old) C57BL6/J male and female mice were obtained from the Jackson Laboratory (JAX) strain (#000664). All animal experiments were conducted according to institutional animal care and safety guidelines at Brigham and Women’s Hospital.

### RNAscope in situ hybridization (mouse)

Mice were euthanized by CO_2_ axphixiation and decapitated. For DRG collection, spines were removed and a saggital cut was used to open up the spine. After the spinal cord was removed, all ipsilateral lumbar ganglia were collected, stored in OCT and frozen in −80 °C. For TG collection, TGs were dissected from the base of the skull after removing the brain. The V1-3 and proximal projections were severed and TGs were stored in OCT and frozen in −80 °C.

Mouse RNAscope florescence in situ hybridization (FISH) experiments were performed according to the manufacturer’s instructions by the Harvard Neurobiology Imaging Facility using the RNAscope Flurescent Multiplex kit (Advanced Cell Diagnostics (ACD)) for fresh frozen tissue. Briefly, mice were anesthetized using isofluorane and decapitated. DRGs and TGs were collected, frozen in OCT and sectioned into 12-14 um sections using a cryostat. RNAscope probes against the following genes were ordered from ACD bio: Scn7a (Cat: 548561), Gpr26 (Cat: 317381), Trpm8 (Cat: 420451-C2) and Calca (Cat: 578771-C3). Following FISH, sections were imaged using a 20x objective lens on a Zeiss LSM710 confocal microscope.

### RNAscope in situ hybridization (human)

Human TGs were obtained from consented donors using a rapid autopsy protocol at Mass General Brigham (IRB# 2017P000757). After removal of the brain for neuropathological analysis, Meckle’s cave was identified in the base of the skull and dissected by JKL. After visualizing the ganglia, the V1-3 nerve braches and the cranial nerve 5 root were cut as they emerged from the ganglion. The dissected ganglia were flash-frozen in liquid nitrogen, and subsequently stored at - 80 °C. The date and time of death, dissection and storage were recorded by pathology staff and donor information was anonymized for downstream processing.

RNAscope fluorescence in situ hybridization (FISH) expereiements were performed according to the manufacturer’s instructions using the RNAscope Fluorescent Multiplex kit (ACD) for fresh froze tissue, as previous described^116^. Briefly, human TG was frozen in OCT and sectioned into 15 μm sections using a cryostat. The following ACDBio RNAscope probes were used: TRPM8-C3 (Cat: 543121-C3) and GPR26-C2 (Cat: 1225951-C2). Channel 1 was left empty to use for lipofuscin subtraction.

### FISH quantification

In mice, three non-consecutive sections from three different animals per probe set were stained and used for quantification. Regions of interests (ROI) that showed puncta in any three probes of the probe channels were manually segmented using imageJ. An ROI was identified as positive for both Gpr26 and Trpm8 if both probes showed 5 puncta. In humans, ROIs were identified as positive if both GPR26 and TRPM8 puncta. We used the green channel (no probe) to detect for lipofuscin and subtracted the signal for the channels we had probes for (GPR26 and TPRM8). Lipofuscin ROI was identified if an ROI had large globular structures in the same pattern across all three channels.

### Human DRG data collection and analysis (HMS)

Human DRGs were obtained from consented organ donors using from the Dr. Gereau’s laboratory at WashU as described below. L4, L5 and T1 DRGs were collected (Table S10).

Single-nuclei of human DRGs were collected using a previously described gradient protocol at HMS^117^. Briefly, human DRGs were initially pulverized on dry ice and approximately 0.5-1 cm^3^ of powder was placed into homogenization buffer (0.25 M sucrose, 25 mM Kcl, 5 mL MgCl2, 20 mM tricine-KOH, pH 7.8, 1 mM DTT, 5 ug/mL actinomycin, 0.04% BSA, and 0.1 U/ul RNase inhibitor) for ∼15s on ice. After the brief incubation for 15s on ice, samples were transferred to a Dounce homogenizer for an additional 10 strokes with a tight pestle in a total volume of 5 mL homogenization buffer. IGEPAL was added to a final concentration of 0.32% and five additional strokes were performed with the tight pestle. The tissue homogenate was then passed through a 40 um filter, and diluted 1:1 with a working solution. Nuclei extracted were layered onto an iodixanol gradient after homogenization and ultracentrifuged as previously described^117^. After ultracentrifugation, nuclei were collected between the 30 and 40% iodixanol layers and diluted for microfluidic encapsulation of individual nuclei in barcoded droplets.

Nuclei suspensions were sequenced using 10X Genomics assays, resuspended and loaded into 10X Chromium device for snRNA-seq (10X Genomics V3.1). Libraries were prepared for snRNA-seq according to the manufacturer’s protocol. Libraries were sequenced in on an Illumina NextSeq 500. Sequencing data were processed and mapped to the human (GRCh38) genome using 10X Genomics Cellranger v7.

### Human DRG data collection (WashU)

Single-nuclei of human DRGs were collected using the gradient protocol mentioned above with the following modifications (Table S10)^117^. Human DRGs were snap-frozen and cut into 100 um sections on a cryostat. Tissue sections were placed into homogenization buffer and after the brief incubation for 15 s on ice, samples were transferred to a 2 mL Dounce homogenizer for 10 strokes with a tight pestle in a total volume of 1 mL homogenization buffer. TritonX-100 was added to a final concentration of 0.1% and five additional strokes were performed with the tight pestle. The rest of the process followed the ordinal protocol previously described^117^.

Nuclei suspensions were loaded into 10X Chromium device for snRNA-seq (10X Genomics V3.1). Libraries were prepared for snRNA-seq according to the manufacturer’s protocol. Libraries were sequenced in on an Illumina NovaSeq 6000 machine. Sequencing data were processed and mapped to the human (GRCh38) genome using 10X Genomics Cellranger v7.

### Human DRG data collection (UTD)

All human tissue procurement procedures were approved by the Institution Review Board at the University of Texas at Dallas (UTD). Lumbar DRGs were recovered from 4 organ donors (2 males, 2 females) age ranging from 32-50 years, and were randomly selected for single nuclei RNA sequencing (Table S10). The tissues were transported from the Southwest Transplant Alliance facility in bubbled aCSF and brought back to the lab for further cleaning to remove excessive connecting tissue and nerves. The trimmed tissue was immediately frozen in pulverized dry ice and stored at −80°C. The frozen tissue was chopped using scissors into small pieces (∼1mm in size) in chilled nuclei isolation buffer (0.25M sucrose, 150mM KCl, 5mM MgCl_2_, 1M Tris Buffer pH 8.0, 0.1mM DTT, protease inhibitor, 0.1% Triton X-100, 0.2U/µl Rnase inhibitor) and then transferred into a Dounce homogenizer containing chilled nuclei isolation buffer. Approximately 15 strokes using the pestle were performed while keeping on ice. The dissociated homogenate was then passed through a 100µm cell strainer and centrifuged using a swing bucket rotor at 500 xg for 10 min at 4°C. The nuclei pellet was resuspended in 2ml of resuspension buffer (1% BSA in PBS, 0.2U/µl Rnase inhibitor) and centrifuged at 500 xg for 5 min at 4°C. The supernatant was removed, and the nuclei pellet was immediately fixed using the 10X Chromium Fixed RNA Profiling kit. The samples were fixed for 16h at 4°C followed by incubation with the 10X Fixed RNA Feature Barcode kit for 16h. The remainder of the library preparation was done according to the manufacturer’s protocol. The samples were sequenced using NextSeq500 at the genome core at the University of Texas at Dallas. Sequencing data were processed and mapped to the human (GRCh38) genome using 10X Genomics Cellranger v7.

### New human DRG data analysis from HMS, UTD, and WashU

Nuclei with > 1000 genes and < 10% of mitochondrial counts were included for analysis. Seurat package (v4) was used to integrate libraries and cluster these nuclei as described by Stuart and colleagues^118^. Briefly, 2000 variable genes were found for each library after normalization and then integration anchors were selected across all libraries. The data were then scaled, and 30 PCs were selected. The libraries were then integrated using Seurat’s CCA algorithm and nuclei were clustered and plotted in UMAP space at a resolution of 1.5. We annotated clusters that were SNAP25+ (log_2_FC > 0.5 and adjusted p.value <0.05 relative to all other clusters) as “Neurons” and clusters that were SPARC+ (log_2_FC > 0.5 and adjusted p.value relative to all other clusters) as “Non-neurons”. At this point, human gene names were converted to mouse gene names using Ensembl’s Biomart ^111^.

Neurons and non-neurons were then separated and reintegrated. Neurons and non-neurons were annotated using marker gene expression (Table S2). Clusters were labelled as doublets if they contained both expression of Snap25, Mpz and Sparc (log_2_FC > 0.5 and adjusted p.value <0.05 relative to all other clusters). Doublets were then removed and the nuclei were reintegrated, and then reannotated.

We then anchored the new human nuclei to the subset of the harmonized atlas comprised of only primate cells/nuclei using Seurat v4 label transfer feature. FindTransferAnchors(reduction = ‘‘cca’’) in Seurat was then used to identify anchors between the new human nuclei and the DRG atlas data. TransferData() was used to transfer atlas subtype labels to each nucleus in the new human data. New human nuclei with anchoring prediction score < 0.5 were excluded from the dataset. Variable genes were identified from the merged dataset, and PCA and UMAP were run to generate new UMAP coordinates.

### Cell type association with UK Biobank GWAS

MAGMA_Celltyping R package (version 1.0.0) was used to calculate cell-type-specific enrichment P values^119^. First, GWAS summary statistics were obtained from UK biobank: Widespread pain for 3+ months (UK Biobank field 2956; 7,575 cases), Pain type(s) experienced in last month: Back pain (field 6159_4; 145,904cases), Pain type(s) experienced in last month: Hip pain (field 6159_6; 64,827 cases), Headaches for 3+ months (field 3799; 49637 cases), migraine (23andme; 30,465 cases). All the SNPs were mapped to a gene-level signature using “GRCh37” and gene-level P values were calculated. Second, raw counts table is normalized to 10,000 using NormalizeData from Seurat (only primate cells/nuclei) and the gene expression specificity is ranked into 40 quantiles for each cell type. Finally, the cell type association analysis was calculated using linear mode. The cell type association P-value is corrected using Benjamini-Hochberg. Variants 1,000 bases upstream and downstream of genes were used in our analysis.

### Positive selection analysis

For our positive selection analysis, we downloaded the Selectome v7 database for human genes^120^. To annotate the genes from the atlas, we converted human gene names in Selectome to mouse gene names using Biomart^111^. For both the DRG and TG, human marker genes (log_2_FC>0.5 relative to all other subtypes in the same species) were then annotated on whether the database reported the gene as under positive selection (column “selected” == 1). We then downloaded sc/snRNA-seq Seurat objects for other nervous tissue and removed any clusters the original studies had annotated as doublets^109,121,122^. Afterwards, FindAllMarkers() was run on each dataset and the markers (log_2_FC>0.5 relative to all other subtypes in the same dataset) were annotated based on whether the Selectome database (human gene names) reported the gene as evolving under positive selection. Amygdala, hippocampus and nucleus accumbens data was downloaded from Tran et al., 2021, motor cortex data was taken from Bakken et al., 2021 and the medial temporal gyrus data was downloaded from the Allen Brain Atlas.

### Analysis of FlyCell atlas data

Loom files for the Drosophila leg (10X Genomics dataset) were downloaded FlyCell using the atlas browser ^90^. The loom files were converted into Seurat objects and then cells were subsetted based on whether they were annotated as “mechanosensory neuron” or “mechanosensory neuron of chordotonal organ” in the metadata’s annotation column. Drosophila gene names were converted to mouse gene names using Diopt v8.5^123^. We then anchored the Drosophila cells to the DRG neuronal reference atlas using Seurat’s label transfer. FindTransferAnchors(reduction = ‘‘cca’’) in Seurat was then used to identify anchors between the DRG atlas and fly data. TransferData() was used to transfer DRG subtype labels to each nucleus/cell in the fly data. Fly cells with anchoring prediction score < 0.5 were excluded from the dataset. Variable genes were identified from the merged dataset, and PCA and UMAP were run to generate new UMAP coordinates.

### Axolotl DRG collection and dissociation at single-cell level

Beta-III tubulin:GAP43-EGFP transgenic axolotls^124^ 7-8cm in length (snout to tail) were anesthetized in 0.1% tricaine. For DRG dissociation at single-cell level, 10X Genomics “Dissociation of Mouse Embryonic Neural Tissue for Single Cell RNA Sequencing” protocol was adapted to axolotl tissue. DRGs of brachial plexus nerves located at C3, C4 and C5 were collected from both left and right sides and combined (total of 6 DRGs) in a single tube containing chilled 0.7X HBSS buffer and kept on ice. For dissociation, 400ul papain (Worthington Biochemical, Cat: #LK003178) dissolved in 0.7X PBS was pre-warmed in 37°C water bath for 10 minutes to activate the enzyme. HBSS was removed and activated papain was added onto DRGs. The DRG in papain was placed in 37°C water bath for 20 minutes. After the incubation, DRGs were settled in the bottom of the tube by a brief spin at 200g for 1min and papain was removed. 500ul DRG media consisting of Neurobasal Plus Medium at 60% (v/v), (Thermo Scientific, cat: A3582901), 50X B-27 Plus Supplement stock at 1X (Thermo Scientific, cat: A3582801), 100X insulin-transferrin-selenium stock at 1% (v/v), (Peprotech, cat: 41400045), 250 μg/mL amphotericin B stock at 1% (v/v), (Sigma-Aldrich, cat: A2942), gentamicin at 50mg/ml (Sigma-Aldrich, cat: G1264), 50ug/ml recombinant human β-NGF stock at 1:1000, (Peprotech, cat: 450-01) mixed in Ringer’s solution (115mM NaCl, 2.5mM KCl, 2mM CaCl2, 10mM HEPES pH7.4, 0.5 mM EDTA dissolved in water and ph adjusted to 7.4 with NaOH) was added. DRGs were triturated in media using 1000 μl pipette. Trituration was performed slowly and gently until most of the tissue was dissociated. After dissociation, sample was centrifuged at 200g for 3min. The supernatant was removed, and the pellet was resuspended in 100 μl of 0.7XPBS +%0.04 BSA. Cell count and single-cell sequencing were performed by Harvard University Bauer Sequencing core. Luna-FL Dual Fluorescence Cell Counter was used to assess number of viable cells. Cell count was repeated 4 times and averaged to determine the final cell count, viability, and cell size. Total of 405 cells/μl with 100% cell viability were present in the final cell suspension in first sample and 486 cells/μl with 97% viability in the second set. scRNA-seq libraries were prepared with the Chromium™ Single Cell 3′ NextGEM, Library and Gel Bead Kit v3.1. Library was sequenced on a NovaSeq 6000 instrument (Illumina) to a depth of 360 million reads with an estimated 45,000 reads per cell. A paired-end reading with 75bp read length was aimed with 28 bases for read 1 and 91 bases for read 2 with 8 bases for index.

### Computational analysis of Axolotl scRNA-seq data

For alignment of 10X reads, we reformatted an axolotl genome^125^ using Cellranger v7. Sequencing data were then mapped to the reformatted axolotl genome using Cellranger v7.

Cells with > 1000 genes and < 10% of mitochondrial counts were included for analysis. Seurat package (v4) was used to integrate libraries from different animals and cluster these nuclei as described by Stuart and colleagues^118^. Briefly, 2000 variable genes were found for each library after normalization and then integration anchors were selected across all libraries. The data were then scaled, and 30 PCs were selected. The libraries were then integrated using Seurat’s CCA algorithm and nuclei were clustered and plotted in UMAP space at a resolution of 1.5. We annotated clusters that were Snap25+ (log_2_FC > 0.5 relative to all other clusters) as “Neurons” and clusters that were Sparc+ (log_2_FC > 0.5 relative to all other clusters) as “Non-neurons”. At this point, axolotl gene names were converted to mouse gene names using Ensembl’s Biomart^111^.

Neurons and non-neurons were then separated and reintegrated. Neurons and non-neurons were annotated using marker gene expression (Table S2). Clusters were labelled as doublets if they contained both expression of Snap25, Mpz and Sparc (log_2_FC > 0.5 relative to all other clusters). Doublets were then removed and the cells were reintegrated.

We labeled clusters enriched for Nefh and Ntrk2 as NF-Ntrk2, clusters enriched for Nefh and Ntrk3 as NF-Ntrk3, and clusters Nefh with no Ntrk2 or Ntrk3 enrichment as NF-unknown. For clusters without for NEFH enrichment, we labelled as CX-1 if we observed Trpm8 enrichment, and CX-2 if we observed Adcyap1 enrichment without Trpm8 enrichment (Table S2). Enrichment was defined as log_2_FC >0.05 relative to all neuronal cluster with an adjusted p.value <0.05. To determine co-expression of Trpv1 and Trpm8, we identified subsetted for cells with greater than 1 count of either Trpv1 or Trpm8. To compare to our mammalian atlas, we subsetted for cells/nuclei from studies that used the 10x Genomics sequencing kit, and then searched for cells with Trpv1 or Trpm8 expression (count >1).

The harmonized DRG atlas was subsetted to only features with 1:1 orthologs in both the DRG atlas and the axolotl data. We then anchored the axolotl cells to the harmonized atlas using the Seurat’s label transfer feature. FindTransferAnchors(reduction = ‘‘cca’’) in Seurat was then used to identify anchors between DRG and TG data. TransferData() was used to transfer DRG subtype labels to each nucleus/cell in the DRG atlas. Axolotl DRG cells with anchoring prediction score < 0.5 were excluded from the dataset. Variable genes were identified from the merged dataset, and PCA and UMAP were run to generate new UMAP coordinates.

### Hybridization Chain Reaction Fluorescence In-Situ Hybridization (HCR-FISH) of sectioned axolotl DRG tissues

Axolotl transcripts used to generate probe sequences were, AMEX60DD055485 for TRPM8, AMEX60DD054745 for TRPV1, and AMEX60DD039896 for TRPA1. A probe generator tool designed by Lovely et al was used to design unique pool of probes to target these genes^126^. Candidate probe sequences were ordered from IDT DNA oPool (Table S11).

Cryopreserved DRG tissue sections collected from naïve animals were stained with all three probes at the same time with the following protocol adapted from Lovely et al^126^. Frozen DRG tissue sections were brought to room temperature, thawed and washed with 2X SSC buffer for 3 x 5min. Tissues were cleared with tissue clearing solution consisting of 4% SDS and 200mM prepared in DEPC-water at pH 8.5. Sections were washed again with 2x SSC 3x 5min. After rinsing sections were incubated in Hybridization buffer (provided by Molecular Instruments https://www.molecularinstruments.com/) for 15min at 37°C.

Probes received from IDT were resuspended in 50ul of TE buffer at 50pmol concentration. Probes were mixed in hybridization buffer at 1:200 dilution and pre-heated at 37°C. After pre-hybridization 100ul probe solution was added onto each slide and covered with parafilm and incubated in 37°C oven overnight. Next day, slides were washed with pre-heated wash buffer (provided by Molecular Instruments) at 37°C 3 × 15min. Slides were then rinsed with 5x SSCT at 37°C for 15min and 5min at room temperature. Excess solutions were removed and 200ul Amplification buffer (provided by Molecular Instruments) was added and slides were incubated at room temperature for 30min. Hairpin solutions (1:50 dilution) were prepared by heating 2ul of 3uM of H1 and H2 hairpins specific for each gene with specific fluorophore (B1 for TRPV1 (647), B2 for TRPM8 (488), B3 for TRPA1 (594)) in separate tubes at 95°C for 90 seconds and cooled to room temperature for about 30min before mixing in 100ul amplification buffer. Hairpin solutions mixed in amplification buffer were applied onto slides, covered with parafilm and incubated at room temperature overnight in dark. On the final day, slides were washed with 5X SSCT 2×30min at room temperature. 200 μl of DAPI was applied for 5min and rinsed with PBS for 5min. Excess solution was removed, and mounting media was added. Stained tissue sections were imaged using Zeiss LSM900 with 40x/1.4N.A oil objective.

## Supplementary Figures

**Figure S1:**
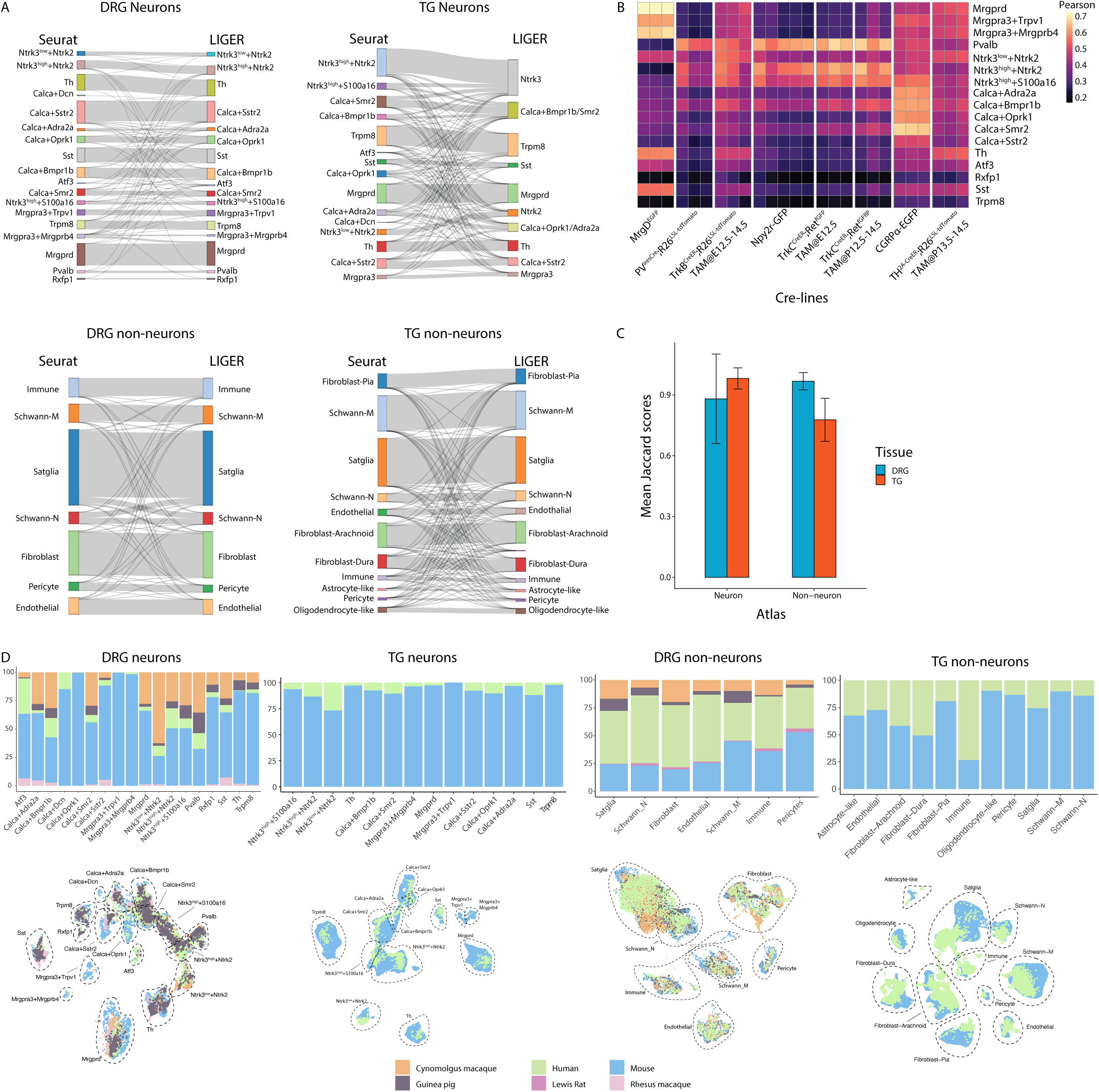
Cell types of the harmonized DRG and TG atlases. *A. Comparison of cell type annotations between Seurat and LIGER.* Sankey plots showing cell type annotations of DRG neurons (top left), TG neurons (top right), DRG non-neurons (bottom left) and TG non-neurons (bottom right) made using Seurat or LIGER pipielines (see Methods). *B. Transcriptomic correlation between atlas cell types and bulk RNA-seq of genetically labeled DRG neuronal populations isolated from Cre lines*. Heatmap displays Pearson’s correlations between the log-normalized counts of the DRG neuronal atlas and the RPKM normalized counts of bulk RNA-seq of DRG neuronal populations isolated Cre lines (3-5 biological replicates)^16^. *C. Jaccard similarity scores for neuronal and non-neuronal cell types*. Displayed are the average Jaccard scores for a subset of neuronal subtypes and non-neuronal subtypes that are present across across all studies (see Methods). Error bars represent standard deviation. *D. Distribution of DRG and TG cell types across species.* Bar plots display the fraction of cells/nuclei from a specific species per cell type in each harmonized atlas. Below each bar plot are the UMAP projections for each atlas colored by species. Each species was downsampled to 3000 cells/nuclei for visualization purposes in the DRG neuronal and non-neuronal atlases.

**Figure S2:**
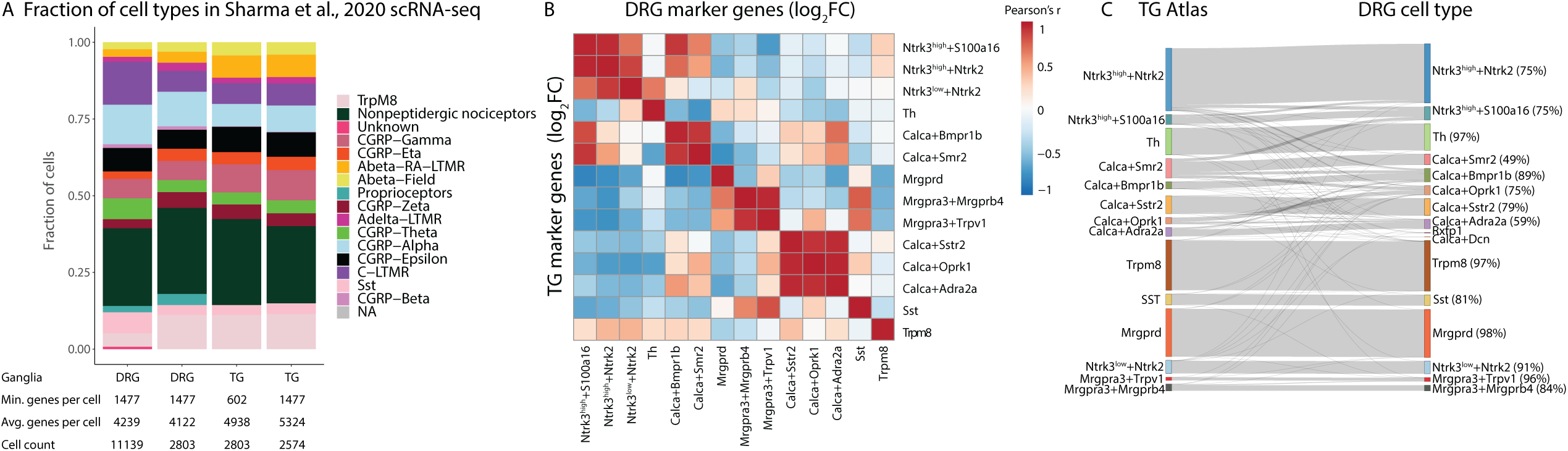
Transcriptomic similarities between TG and DRG neurons. A. *Fraction of cell types in DRG scRNA-seq data from Sharma et al., 2020*. Stacked bar plot shows the fraction of DRG or TG neuronal subtypes from Sharma et al., 2020. The ratio of cell types are also shown when all DRG neurons are downsampled to match the total number of TG neurons, as well as when TG cells were filtered to have the same minimum number of genes per cell as the DRG data (1477 genes). Table below bars summarizes the minimum number of genes per cell (“Min. genes per cell”), mean number of genes per cell (“Avg. genes per cell”) and total number of cells (“Cell count”) for each bar. B. *Comparison of cell-type-specific gene expression patterns in the DRG and TG*. Displayed is the Pearson correlation of marker gene log_2_FC values between each TG and DRG cell type (log_2_FC > 1, adjusted p-value <0.05 relative to all neuronal cells in the same ganglia). C. *Comparison of TG cells/nuclei annotation by anchoring to DRG atlas*. Gray lines connect the the TG harmonized atlas cell type annotations (left) and the DRG cell to which they are most transcriptomically similar when anchored to the harmonized DRG atlas (right). The percentage of TG cells/nuclei that anchored to the corresponding DRG cell type are shown. Only cells/nuclei with anchoring prediction score >0.5 are shown.

**Figure S3:**
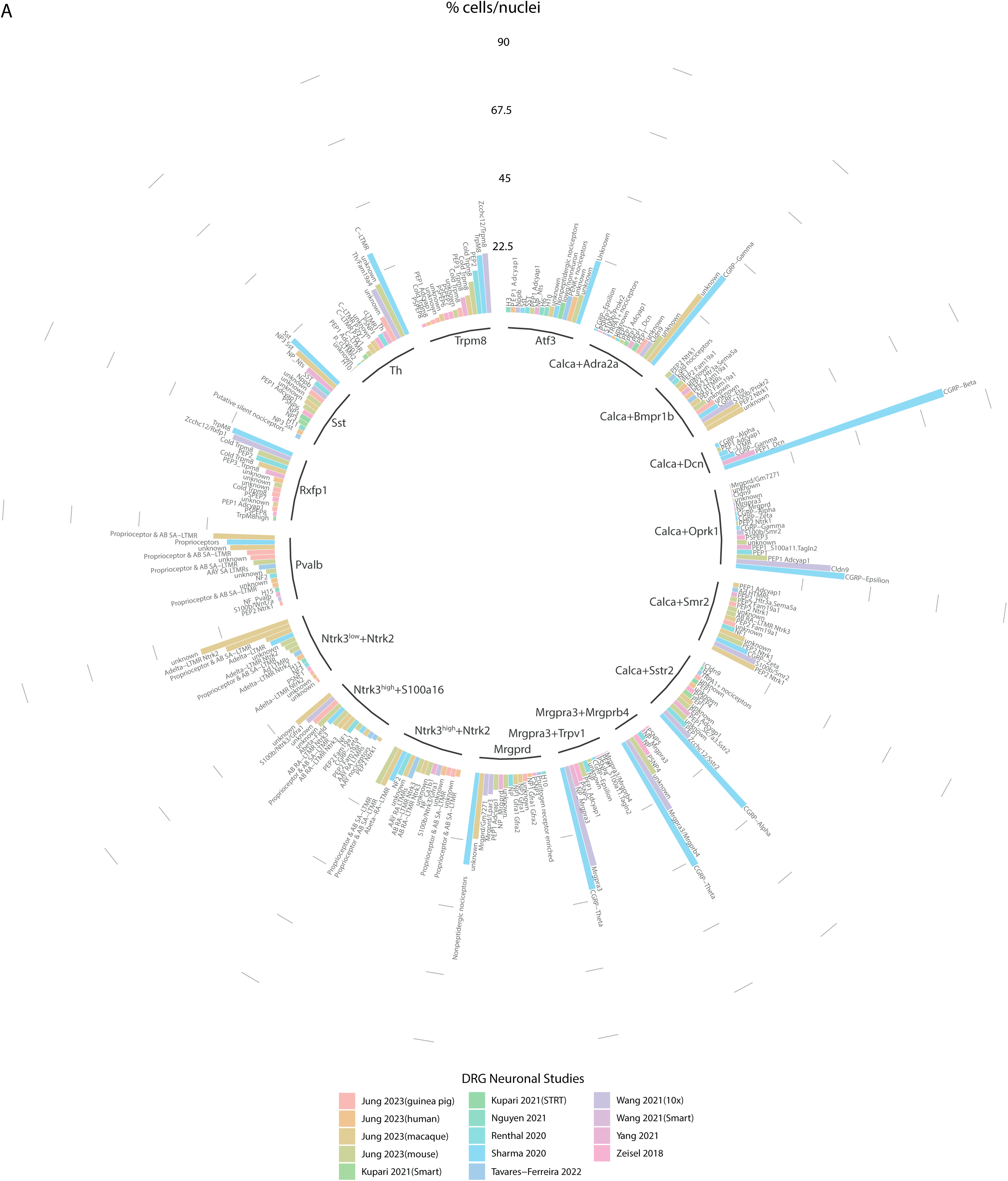
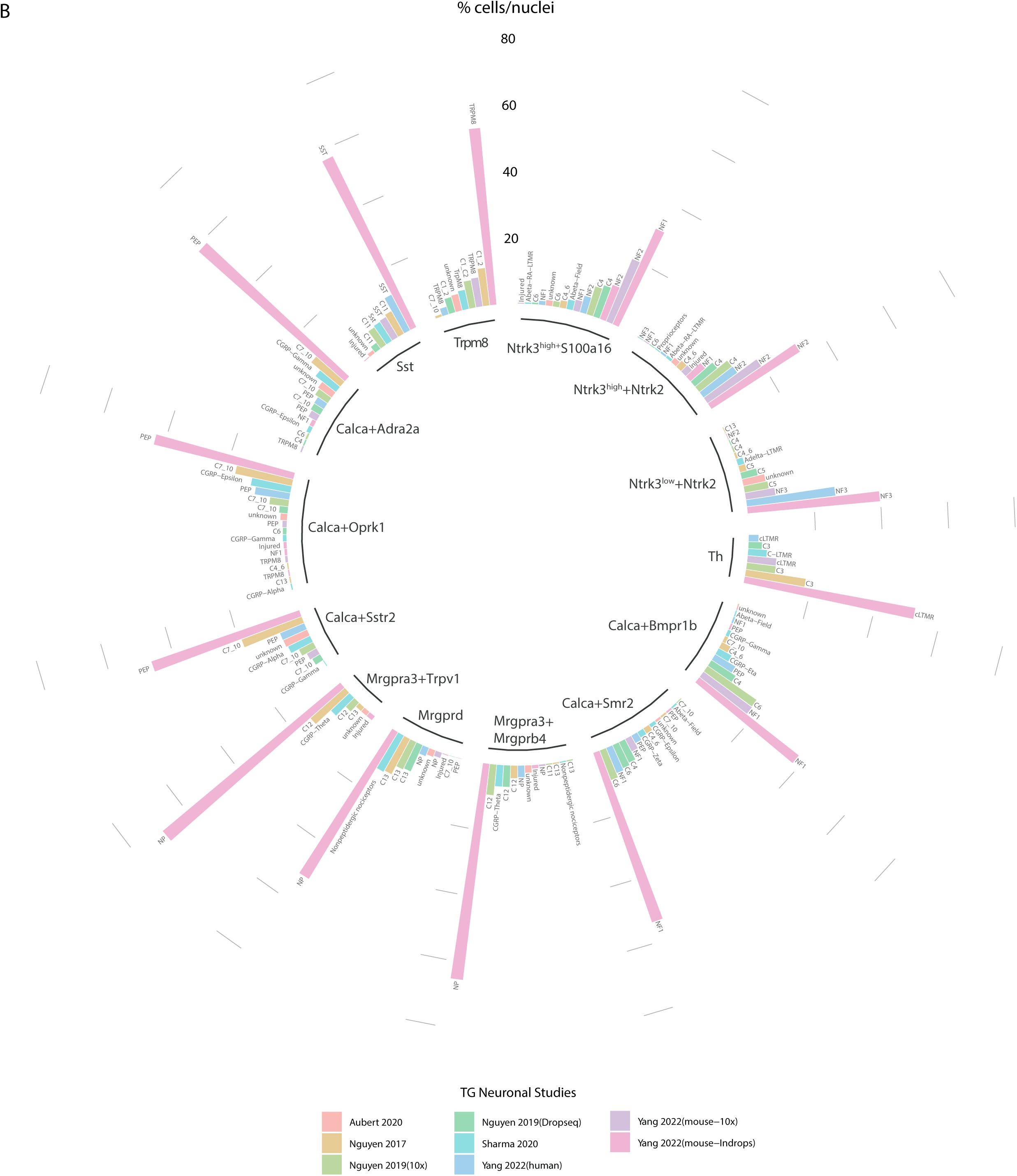
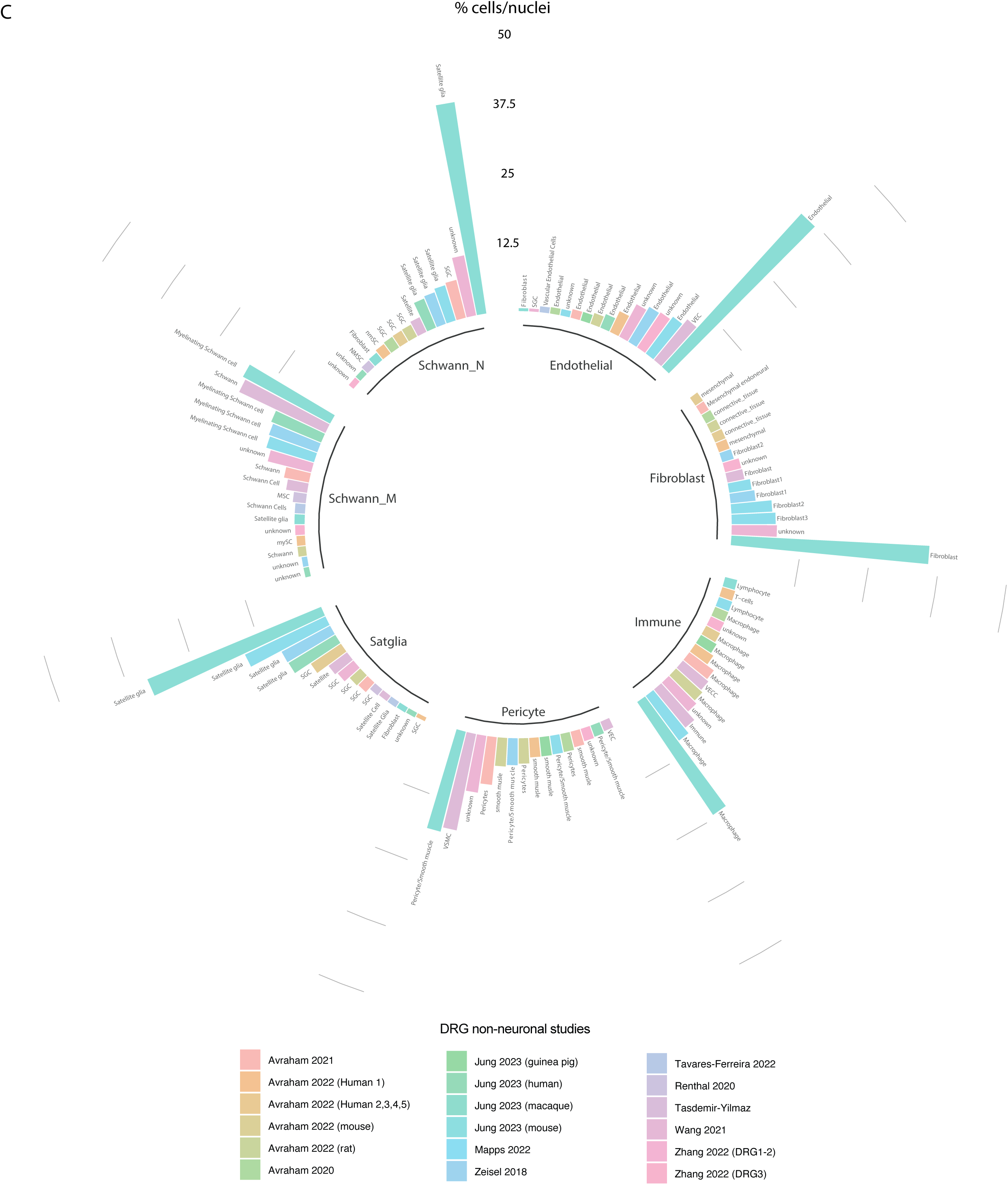
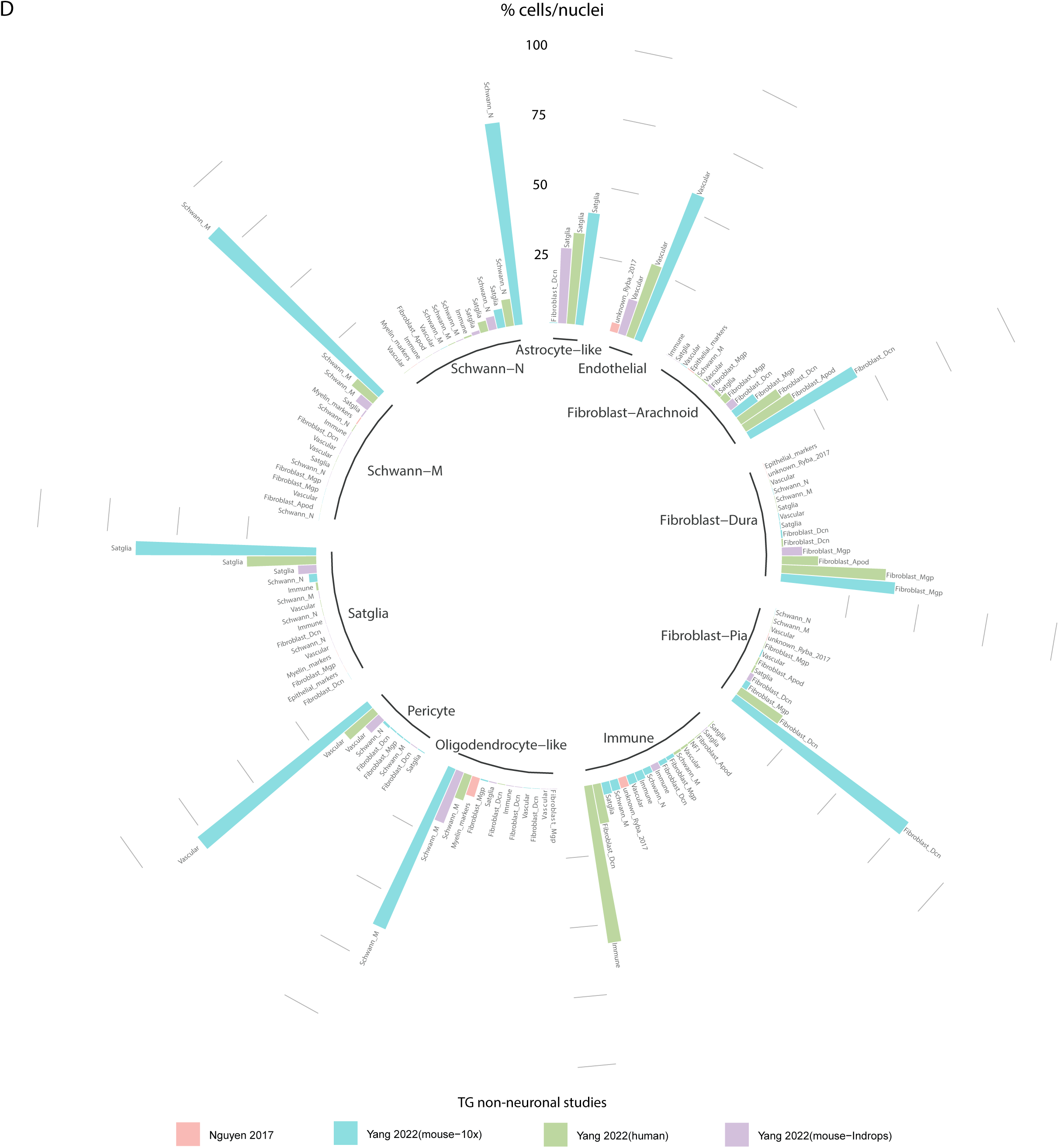
Comparison of nomenclature between DRG or TG atlases and the individual sc/snRNA-seq studies that comprise them. Circle bar plots display the fraction of cells/nuclei from each individual DRG or TG sc/snRNA-seq study that contributes to the harmonized cell type in the *DRG neuronal atlas (A), TG neuronal atlas (B), DRG non-neuronal atlas (C) and TG non-neuronal atlas (D)*. The harmonized atlas cell type nomenclature is shown inside the circle and the corresponding cell type nomenclature defined in individual sc/snRNA-seq studies is shown on top of its respective bar. Bar color corresponds to the individual DRG or TG sc/snRNA-seq study.

**Figure S4:**
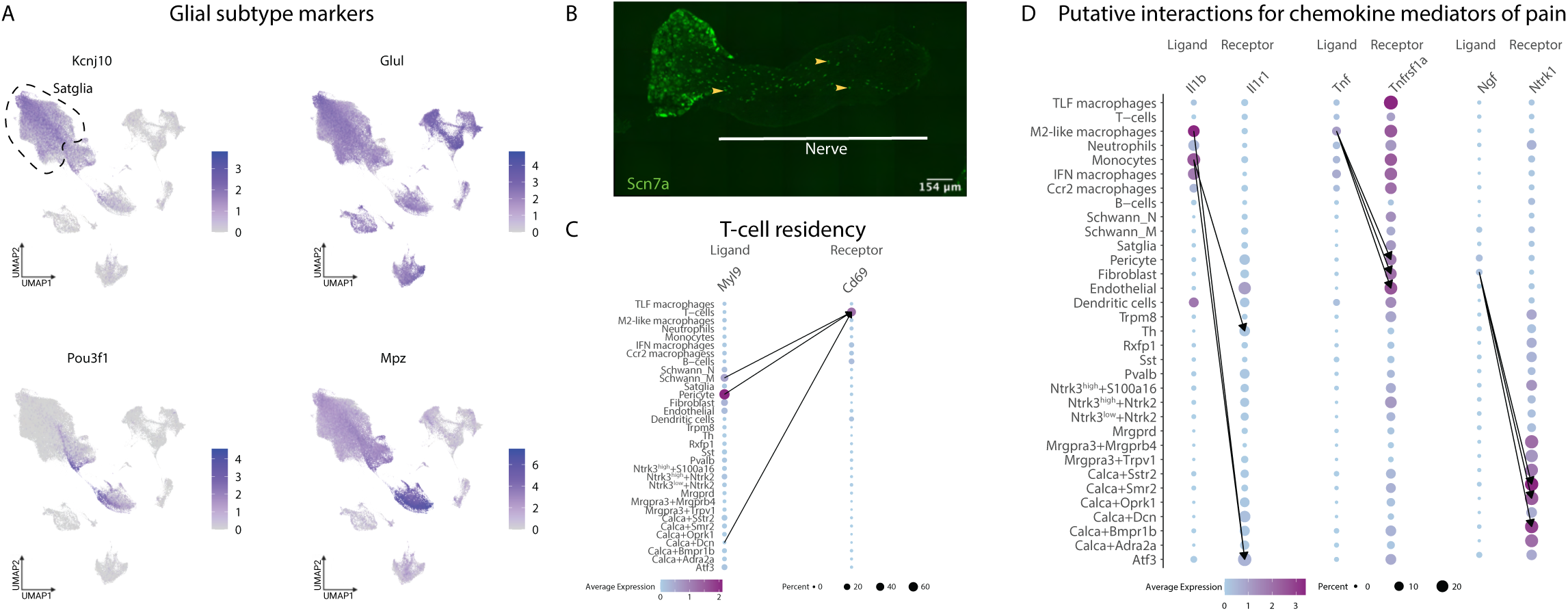
Cell-type-specific gene expression in DRG glial and immune cells. *A. Expression profiles of satellite glia subtype marker genes.* UMAP displays gene expression of satellite glia subtype marker genes across 187,383 cells/nuclei. Cells/nuclei are colored by log-normalized scaled gene expression. *B. Scn7a expression in DRG.* Example florescent *in situ* hybridization of DRG showing expression of *Scn7a* in DRG. Arrows point to *Scn7a* expression in Schwann_N cells present in the nerve. *C. Predicted ligand receptor interaction between Myl9 and Cd69*. Dot plot of ligand or receptor expression in DRG cell types. Dot size indicates the percent of cells/nuclei expressing the ligand or receptor and color indicates average log-normalized gene expression. Arrows connect the cell-cell interactions that have the three highest ligand-receptor scores (see Methods). *D. Predicted ligand receptor interactions of cytokine and chemokine mediators of pain between DRG cell types.* Dot plot of ligand or receptor expression in DRG cell types. Dot size indicates the fraction of cells/nuclei expressing the ligand or receptor and color indicates average log-normalized gene expression. Arrows connect the cell-cell interactions that have the 3 highest ligand-receptor scores (see Methods).

**Figure S5:**
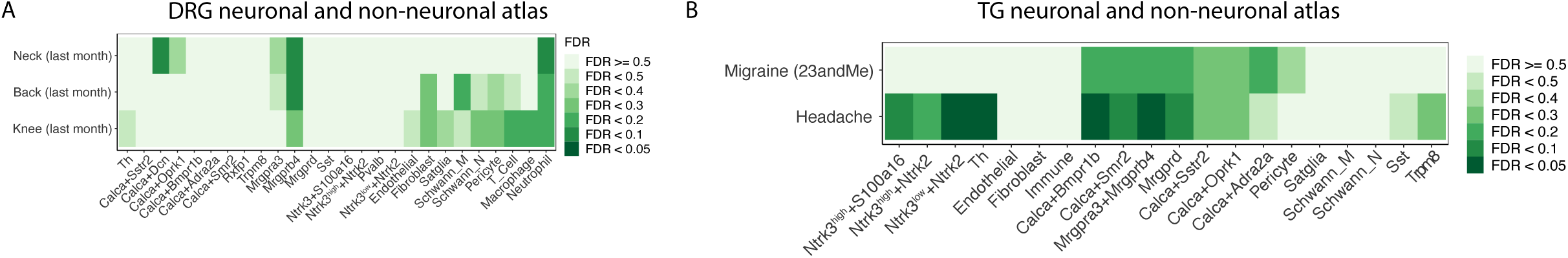
DRG and TG cell type enrichment of genes associated with UK Biobank pain and headache genome-wide association studies. Heatmap displays the Benjamini-Hochberg corrected enrichment of genomic variants associated with pain or headache conditions (UK Biobank or 23andMe) in the genomic regions including genes differentially expressed (see Methods) in each (A) DRG or (B) TG cell types relative to the genomic regions including genes differentially expressed in all other cell types of the same ganglia.

**Figure S6:**
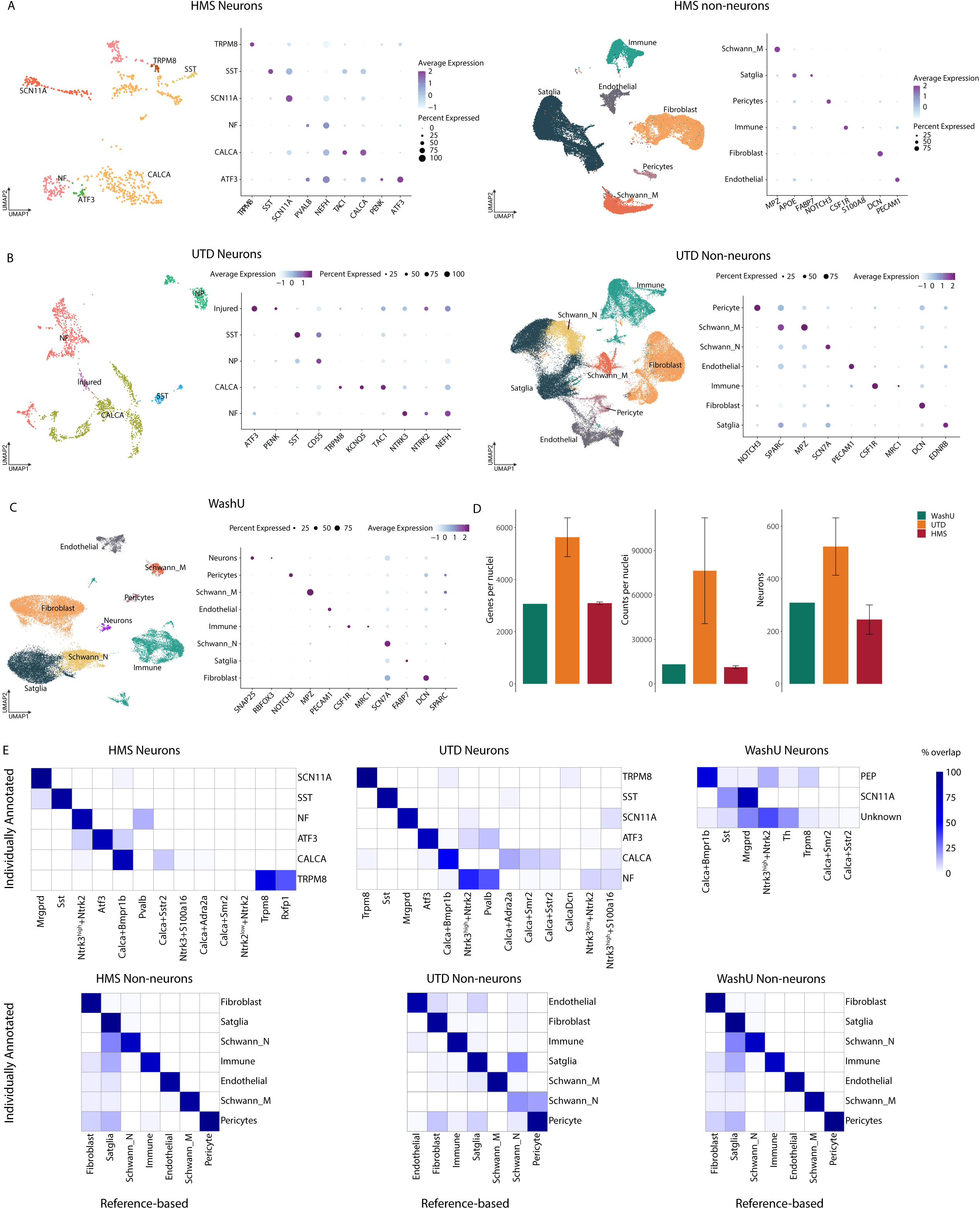
Annotation of new human DRG snRNA-seq data from three institutions. *A. Individual annotation of HMS DRG snRNA-seq data.* Left: UMAP projection of neuronal HMS snRNA-seq data (762 nuclei) and corresponding dot plot of marker gene expression. Right: UMAP projection of non-neuronal HMS snRNA-seq data (39,772 nuclei) and corresponding dot plot of marker gene expression. UMAPs are colored by cell type. Dot size indicates the fraction of cells/nuclei expressing each gene and color indicates average log-normalized scaled expression of each gene. *B*. *Individual annotation of UTD snRNA-seq data*. Left: UMAP projection of neuronal UTD snRNA-seq data (2,614 nuclei) and corresponding dot plot of marker gene expression. Right: UMAP projection of non-neuronal UTD snRNA-seq data (61,336 nuclei) and corresponding dot plot of marker gene exprssion. UMAPs are colored by cell type. Dot size indicates the fraction of cells/nuclei expressing each gene and color indicates average log-normalized scaled expression of each gene. *C*. *Individual annotation of WashU snRNA-seq data.* UMAP projection of all UTD snRNA-seq data (31,350 nuclei) and corresponding dot plot of marker gene expression. UMAP is colored by cell type. Dot size indicates the fraction of cells/nuclei expressing each gene and color indicates average log-normalized scaled expression of each gene. D. *Comparison of HMS, UTD and WashU snRNA-seq data metrics.* Bar plots indicate the number of genes per nucleus, number of unique molecular identifiers (UMI) per nucleus, and the number of neurons per dataset. Error bars represent standard deviation across distinct libraries in each institute’s dataset. *E*. *Comparison of indivudal and reference-based cell type assignment of new human DRG snRNA-seq data*. For each institute, heatmaps display overlap between cell type annotations assigned by analysis indivudally (rows) and the reference-based annotations assigned by anchoring to the harmonized DRG atlas (columns) (see Methods). Color indicates percent overlap.

**Figure S7:**
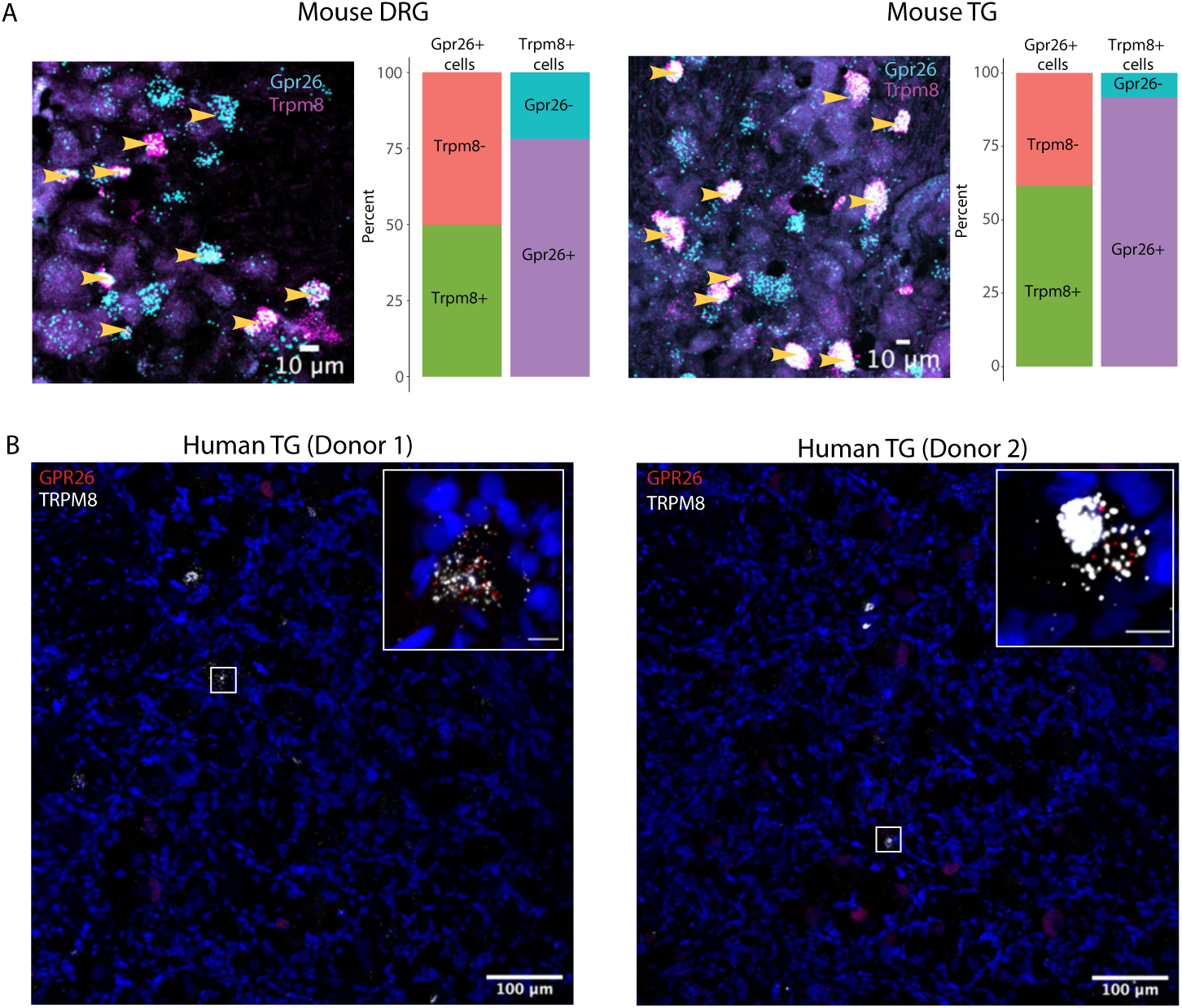
*GPR26* and *TRPM8* expression across species. A. *Co-expression of Gpr26 and Trpm8 mouse DRG and TG.* Examples of *in situ* hybridization of *Grp26 and Trpm8* in DRG and TG and quantification of the fraction of neurons that co-express *Gpr26* and *Trpm8*. Left bar represents the proportion of *Gpr26+* cells that are *Trpm8+*, right bar represents the proportion of *Trpm8+* cells that are *Gpr26+* (DRG n = 36 cells across 3 mice, TG n = 49 cells across 4 mice). B. *Co-expression of GPR26 and TRPM8 in human TG.* Examples of *in situ* hybridization of *GRP26* and *TRPM8* in human TG. Inset is a 7.4x zoom of a *GPR26*+/*TRPM8*+ neuron. Inset scale represents 10 um.

**Figure S8:**
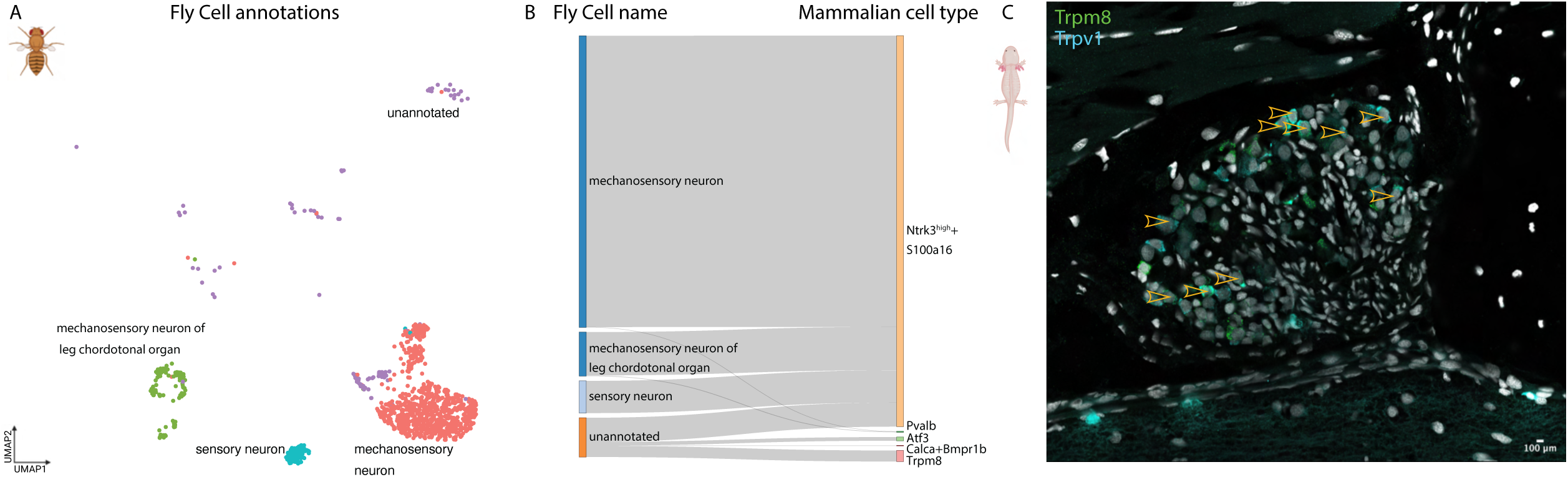
Transcriptomic similarity between non-mammalian and mamammalian peripheral sensory neurons. *A*. *Drosophila leg mechanosensory neurons*. UMAP projection of *Drosophila* leg neurons (n = 2,485 cells) from Fly Cell Atlas. Colors represent cell types. *B*. *Drosophila leg mechanosensory neurons are transcriptomically similar to mammalian Ntrk3^high^+S100a16 neurons*. Gray lines connect the the Fly Cell cell type annotation (left) with the mammalian DRG neuronal subtype to which they are most transcriptomically similar when anchored to the harmonized DRG atlas (right). Only cells/nuclei with anchoring prediction score >0.5 are shown. Over 99% of *Drosphila* mechanosensory cells anchor to mammalian *Ntrk3^high^+S100a16* (Aβ Field-LTMR). In constrast only ∼6% of *Drosophila* mechanosensory cells have anchoring scores > 0.5 when anchored to the DRG non-neuronal atlas (see Methods). C. *Trpm8 and Trpv1 expression in axolotl DRG neurons.* Example image of *Trpm8* and *Trpv1* hybridization chain reaction *in situ* hybridization. Arrows point to cells co-expressing *Trpm8* and *Trpv1*.

## References

1. Abraira, V. E. & Ginty, D. D. The Sensory Neurons of Touch. Neuron 79, 618–639 (2013).

2. Zimmerman, A., Bai, L. & Ginty, D. D. The gentle touch receptors of mammalian skin. Science 346, 950–954 (2014).

3. Emery, E. C. & Ernfors, P. Dorsal Root Ganglion Neuron Types and Their Functional Specialization. in The Oxford Handbook of the Neurobiology of Pain (ed. Wood, J. N.) 0 (Oxford University Press, 2020). doi:10.1093/oxfordhb/9780190860509.013.4.

4. Ma, Q. A functional subdivision within the somatosensory system and its implications for pain research. Neuron 110, 749–769 (2022).

5. Babetto, E., Wong, K. M. & Beirowski, B. A glycolytic shift in Schwann cells supports injured axons. Nat. Neurosci. 23, 1215–1228 (2020).

6. Singhmar, P. et al. The fibroblast-derived protein PI16 controls neuropathic pain. Proc. Natl. Acad. Sci. 117, 5463–5471 (2020).

7. Chen, Z. et al. Purinergic signaling between neurons and satellite glial cells of mouse dorsal root ganglia modulates neuronal excitability in vivo. Pain 163, 1636–1647 (2022).

8. Harper, A. A. & Lawson, S. N. Conduction velocity is related to morphological cell type in rat dorsal root ganglion neurones. J. Physiol. 359, 31–46 (1985).

9. White, D. M. & Levine, J. D. Different mechanical transduction mechanisms for the immediate and delayed responses of rat C-fiber nociceptors. J. Neurophysiol. (1991) doi:10.1152/jn.1991.66.2.363.

10. Li, L. et al. The functional organization of cutaneous low-threshold mechanosensory neurons. Cell 147, 1615–1627 (2011).

11. Kupari, J. & Ernfors, P. Molecular taxonomy of nociceptors and pruriceptors. PAIN 10.1097/j.pain.0000000000002831 (2022) doi:10.1097/j.pain.0000000000002831.

12. Marshall, K. L. et al. PIEZO2 in sensory neurons and urothelial cells coordinates urination. Nature 588, 290–295 (2020).

13. Wang, Y. et al. The role of somatosensory innervation of adipose tissues. Nature 609, 569– 574 (2022).

14. Wolfson, R. L. et al. DRG afferents that mediate physiologic and pathologic mechanosensation from the distal colon. 2022.11.27.518103 Preprint at https://doi.org/10.1101/2022.11.27.518103 (2022).

15. Marmigère, F. & Ernfors, P. Specification and connectivity of neuronal subtypes in the sensory lineage. Nat. Rev. Neurosci. 8, 114–127 (2007).

16. Zheng, Y. et al. Deep Sequencing of Somatosensory Neurons Reveals Molecular Determinants of Intrinsic Physiological Properties. Neuron 103, 598–616.e7 (2019).

17. Zeisel, A. et al. Molecular Architecture of the Mouse Nervous System. Cell 174, 999–1014.e22 (2018).

18. Nguyen, M. Q., Le Pichon, C. E. & Ryba, N. Stereotyped transcriptomic transformation of somatosensory neurons in response to injury. eLife 8, e49679 (2019).

19. Aubert, M. et al. Gene editing and elimination of latent herpes simplex virus in vivo. Nat. Commun. 11, 4148 (2020).

20. Avraham, O. et al. Satellite glial cells promote regenerative growth in sensory neurons. Nat. Commun. 11, 4891 (2020).

21. Avraham, O. et al. Profiling sensory neuron microenvironment after peripheral and central axon injury reveals key pathways for neural repair. eLife 10, e68457 (2021).

22. Avraham, O. et al. Profiling the molecular signature of satellite glial cells at the single cell level reveals high similarities between rodents and humans. Pain 163, 2348–2364 (2022).

23. Sharma, N. et al. The emergence of transcriptional identity in somatosensory neurons. Nature 577, 392–398 (2020).

24. Kupari, J. et al. Single cell transcriptomics of primate sensory neurons identifies cell types associated with chronic pain. Nat. Commun. 12, 1510 (2021).

25. Nguyen, M. Q., von Buchholtz, L. J., Reker, A. N., Ryba, N. J. & Davidson, S. Single-nucleus transcriptomic analysis of human dorsal root ganglion neurons. eLife 10, e71752 (2021).

26. Wang, K. et al. Single-cell transcriptomic analysis of somatosensory neurons uncovers temporal development of neuropathic pain. Cell Res. 31, 904–918 (2021).

27. Mapps, A. A. et al. Diversity of satellite glia in sympathetic and sensory ganglia. Cell Rep. 38, 110328 (2022).

28. Tavares-Ferreira, D. et al. Spatial transcriptomics of dorsal root ganglia identifies molecular signatures of human nociceptors. Sci. Transl. Med. 14, eabj8186 (2022).

29. Yang, L. et al. Human and mouse trigeminal ganglia cell atlas implicates multiple cell types in migraine. Neuron (2022) doi:10.1016/j.neuron.2022.03.003.

30. Zhang, D. et al. Single-nucleus transcriptomic analysis reveals divergence of glial cells in peripheral somatosensory system between human and mouse. 2022.02.15.480622 Preprint at https://doi.org/10.1101/2022.02.15.480622 (2022).

31. Jung, M. et al. Cross-species transcriptomic atlas of dorsal root ganglia reveals species-specific programs for sensory function. Nat. Commun. 14, 366 (2023).

32. Qi, L. et al. A DRG genetic toolkit reveals molecular, morphological, and functional diversity of somatosensory neuron subtypes. 2023.04.22.537932 Preprint at https://doi.org/10.1101/2023.04.22.537932 (2023).

33. Tran, H. T. N. et al. A benchmark of batch-effect correction methods for single-cell RNA sequencing data. Genome Biol. 21, 12 (2020).

34. Luecken, M. D. et al. Benchmarking atlas-level data integration in single-cell genomics. Nat. Methods 19, 41–50 (2022).

35. Yu, H. et al. Single-Soma Deep RNA sequencing of Human DRG Neurons Reveals Novel Molecular and Cellular Mechanisms Underlying Somatosensation. 2023.03.17.533207 Preprint at https://doi.org/10.1101/2023.03.17.533207 (2023).

36. Butler, A., Hoffman, P., Smibert, P., Papalexi, E. & Satija, R. Integrating single-cell transcriptomic data across different conditions, technologies, and species. Nat. Biotechnol. (2018) doi:10.1038/nbt.4096.

37. Welch, J. D. et al. Single-Cell Multi-omic Integration Compares and Contrasts Features of Brain Cell Identity. Cell 177, 1873–1887.e17 (2019).

38. Russ, D. E. et al. A harmonized atlas of mouse spinal cord cell types and their spatial organization. Nat. Commun. 12, 5722 (2021).

39. Ernfors, P., Lee, K. F., Kucera, J. & Jaenisch, R. Lack of neurotrophin-3 leads to deficiencies in the peripheral nervous system and loss of limb proprioceptive afferents. Cell 77, 503–512 (1994).

40. Arber, S., Ladle, D. R., Lin, J. H., Frank, E. & Jessell, T. M. ETS Gene Er81 Controls the Formation of Functional Connections between Group Ia Sensory Afferents and Motor Neurons. Cell 101, 485–498 (2000).

41. Jc, de N., S, D. & Tm, J. Etv1 inactivation reveals proprioceptor subclasses that reflect the level of NT3 expression in muscle targets. Neuron 77, (2013).

42. Luo, W., Enomoto, H., Rice, F. L., Milbrandt, J. & Ginty, D. D. Molecular Identification of Rapidly Adapting Mechanoreceptors and their Developmental Dependence on Ret Signaling. Neuron 64, 841–856 (2009).

43. Bai, L. et al. Genetic identification of an expansive mechanoreceptor sensitive to skin stroking. Cell 163, 1783–1795 (2015).

44. Rutlin, M. et al. The Cellular and Molecular Basis of Direction Selectivity of Aδ-LTMRs. Cell 159, 1640–1651 (2014).

45. McKemy, D. D., Neuhausser, W. M. & Julius, D. Identification of a cold receptor reveals a general role for TRP channels in thermosensation. Nature 416, 52–58 (2002).

46. Bautista, D. M. et al. The menthol receptor TRPM8 is the principal detector of environmental cold. Nature 448, 204–208 (2007).

47. Dhaka, A. et al. TRPM8 Is Required for Cold Sensation in Mice. Neuron 54, 371–378 (2007).

48. Yarmolinsky, D. A. et al. Coding and plasticity in the mammalian thermosensory system. Neuron 92, 1079–1092 (2016).

49. Lou, S., Duan, B., Vong, L., Lowell, B. B. & Ma, Q. Runx1 Controls Terminal Morphology and Mechanosensitivity of VGLUT3-expressing C-Mechanoreceptors. J. Neurosci. 33, 870– 882 (2013).

50. Zylka, M. J., Rice, F. L. & Anderson, D. J. Topographically Distinct Epidermal Nociceptive Circuits Revealed by Axonal Tracers Targeted to Mrgprd. Neuron 45, 17–25 (2005).

51. Stantcheva, K. K. et al. A subpopulation of itch-sensing neurons marked by Ret and somatostatin expression. EMBO Rep. 17, 585–600 (2016).

52. Qi, L. et al. Hierarchical Specification of Pruriceptors by Runt-Domain Transcription Factor Runx1. J. Neurosci. 37, 5549–5561 (2017).

53. Olson, W. et al. Sparse genetic tracing reveals regionally specific functional organization of mammalian nociceptors. eLife 6, e29507 (2017).

54. Elias, L. J. et al. Identification of touch neurons underlying dopaminergic pleasurable touch and sexual receptivity. 2021.09.22.461355 Preprint at https://doi.org/10.1101/2021.09.22.461355 (2022).

55. Tobori, S. et al. MrgprB4 in trigeminal neurons expressing TRPA1 modulates unpleasant sensations. J. Pharmacol. Sci. 146, 200–205 (2021).

56. Vrontou, S., Wong, A. M., Rau, K. K., Koerber, H. R. & Anderson, D. J. Genetic identification of C fibres that detect massage-like stroking of hairy skin in vivo. Nature 493, 669–673 (2013).

57. Lu, P. et al. MrgprA3-expressing pruriceptors drive pruritogen-induced alloknesis through mechanosensitive Piezo2 channel. 2022.06.22.497257 Preprint at https://doi.org/10.1101/2022.06.22.497257 (2022).

58. Han, L. et al. A subpopulation of nociceptors specifically linked to itch. Nat. Neurosci. 16, 174–182 (2013).

59. Hill, R. Z., Loud, M. C., Dubin, A. E., Peet, B. & Patapoutian, A. PIEZO1 transduces mechanical itch in mice. Nature 607, 104–110 (2022).

60. Huang, J. et al. Circuit dissection of the role of somatostatin in itch and pain. Nat. Neurosci. 21, 707–716 (2018).

61. Voisin, T. et al. The CysLT2R receptor mediates leukotriene C4-driven acute and chronic itch. Proc. Natl. Acad. Sci. U. S. A. 118, e2022087118 (2021).

62. Tsujino, H. et al. Activating transcription factor 3 (ATF3) induction by axotomy in sensory and motoneurons: A novel neuronal marker of nerve injury. Mol. Cell. Neurosci. 15, 170– 182 (2000).

63. Parsadanian, A., Pan, Y., Li, W., Myckatyn, T. M. & Brakefield, D. Astrocyte-derived transgene GDNF promotes complete and long-term survival of adult facial motoneurons following avulsion and differentially regulates the expression of transcription factors of AP-1 and ATF/CREB families. Exp. Neurol. 200, 26–37 (2006).

64. Renthal, W. et al. Transcriptional Reprogramming of Distinct Peripheral Sensory Neuron Subtypes after Axonal Injury. Neuron 108, 128–144.e9 (2020).

65. Chu, Y. et al. Single-cell transcriptomic profile of satellite glial cells in trigeminal ganglion. Front. Mol. Neurosci. 16, (2023).

66. Scherer, S. S. et al. Transgenic Expression of Human Connexin32 in Myelinating Schwann Cells Prevents Demyelination in Connexin32-Null Mice. J. Neurosci. 25, 1550–1559 (2005).

67. Watanabe, E., Hiyama, T. Y., Kodama, R. & Noda, M. NaX sodium channel is expressed in non-myelinating Schwann cells and alveolar type II cells in mice. Neurosci. Lett. 330, 109– 113 (2002).

68. Feng, R., Muraleedharan Saraswathy, V., Mokalled, M. H. & Cavalli, V. Self-renewing macrophages in dorsal root ganglia contribute to promote nerve regeneration. Proc. Natl. Acad. Sci. U. S. A. 120, e2215906120 (2023).

69. Lund, H. et al. A network of CD163+ macrophages monitors enhanced permeability at the blood-dorsal root ganglion barrier. 2023.03.27.534318 Preprint at https://doi.org/10.1101/2023.03.27.534318 (2023).

70. Davies, L. C., Jenkins, S. J., Allen, J. E. & Taylor, P. R. Tissue-resident macrophages. Nat. Immunol. 14, 986–995 (2013).

71. Dick, S. A. et al. Self-renewing resident cardiac macrophages limit adverse remodeling following myocardial infarction. Nat. Immunol. 20, 29–39 (2019).

72. Yángüez, E. et al. ISG15 Regulates Peritoneal Macrophages Functionality against Viral Infection. PLoS Pathog. 9, e1003632 (2013).

73. Alivernini, S. et al. Distinct synovial tissue macrophage subsets regulate inflammation and remission in rheumatoid arthritis. Nat. Med. 26, 1295–1306 (2020).

74. Munnur, D. et al. Altered ISGylation drives aberrant macrophage-dependent immune responses during SARS-CoV-2 infection. Nat. Immunol. 22, 1416–1427 (2021).

75. Kurowska-Stolarska, M. & Alivernini, S. Synovial tissue macrophages in joint homeostasis, rheumatoid arthritis and disease remission. Nat. Rev. Rheumatol. 18, 384–397 (2022).

76. Gebhardt, T. et al. Memory T cells in nonlymphoid tissue that provide enhanced local immunity during infection with herpes simplex virus. Nat. Immunol. 10, 524–530 (2009).

77. Szabo, P. A., Miron, M. & Farber, D. L. Location, location, location: Tissue resident memory T cells in mice and humans. Sci. Immunol. 4, eaas9673 (2019).

78. Klein, U., Rajewsky, K. & Küppers, R. Human Immunoglobulin (Ig)M+IgD+ Peripheral Blood B Cells Expressing the CD27 Cell Surface Antigen Carry Somatically Mutated Variable Region Genes: CD27 as a General Marker for Somatically Mutated (Memory) B Cells. J. Exp. Med. 188, 1679–1689 (1998).

79. Allie, S. R. & Randall, T. D. Resident Memory B Cells. Viral Immunol. 33, 282–293 (2020).

80. Kläsener, K. et al. CD20 as a gatekeeper of the resting state of human B cells. Proc. Natl. Acad. Sci. U. S. A. 118, e2021342118 (2021).

81. O’Connell, F. P., Pinkus, J. L. & Pinkus, G. S. CD138 (syndecan-1), a plasma cell marker immunohistochemical profile in hematopoietic and nonhematopoietic neoplasms. Am. J. Clin. Pathol. 121, 254–263 (2004).

82. McCarron, M. J., Park, P. W. & Fooksman, D. R. CD138 mediates selection of mature plasma cells by regulating their survival. Blood 129, 2749–2759 (2017).

83. Burgess, M., Wicks, K., Gardasevic, M. & Mace, K. A. Cx3CR1 Expression Identifies Distinct Macrophage Populations That Contribute Differentially to Inflammation and Repair. ImmunoHorizons 3, 262–273 (2019).

84. Wang, P. L. et al. Peripheral nerve resident macrophages share tissue-specific programming and features of activated microglia. Nat. Commun. 11, 2552 (2020).

85. Kimura, M. Y., Koyama-Nasu, R., Yagi, R. & Nakayama, T. A new therapeutic target: the CD69-Myl9 system in immune responses. Semin. Immunopathol. 41, 349–358 (2019).

86. Hayashizaki, K., et al. Myosin light chains 9 and 12 are functional ligands for CD69 that regulate airway inflammation. Sci. Immunol. 1, eaaf9154 (2016).

87. Yadav, A. et al. A cellular taxonomy of the adult human spinal cord. Neuron 111, 328–344.e7 (2023).

88. Duan, B. et al. Identification of Spinal Circuits Transmitting and Gating Mechanical Pain. Cell 159, 1417–1432 (2014).

89. Gatto, G. et al. A Functional Topographic Map for Spinal Sensorimotor Reflexes. Neuron 109, 91–104.e5 (2021).

90. Li, H. et al. Fly Cell Atlas: A single-nucleus transcriptomic atlas of the adult fruit fly. Science 375, eabk2432.

91. Cao, J. et al. Comprehensive single cell transcriptional profiling of a multicellular organism. Science 357, 661–667 (2017).

92. Leigh, N. D. et al. Transcriptomic landscape of the blastema niche in regenerating adult axolotl limbs at single-cell resolution. Nat. Commun. 9, 5153 (2018).

93. Payzin-Dogru, D. et al. Nerve-mediated amputation-induced stem cell activation primes distant appendages for future regeneration events in axolotl. 2021.12.29.474455 Preprint at https://doi.org/10.1101/2021.12.29.474455 (2021).

94. Oda, M., Ogino, H., Kubo, Y. & Saitoh, O. Functional properties of axolotl transient receptor potential ankyrin 1 revealed by the heterologous expression system. Neuroreport 30, 323– 330 (2019).

95. Hori, S. & Saitoh, O. Unique high sensitivity to heat of axolotl TRPV1 revealed by the heterologous expression system. Biochem. Biophys. Res. Commun. 521, 914–920 (2020).

96. Gracheva, E. O. & Bagriantsev, S. N. Evolutionary adaptation to thermosensation. Curr. Opin. Neurobiol. 34, 67–73 (2015).

97. Key, F. M. et al. Human local adaptation of the TRPM8 cold receptor along a latitudinal cline. PLoS Genet. 14, e1007298 (2018).

98. Tamuri, A. U. & Dos Reis, M. A Mutation-Selection Model of Protein Evolution under Persistent Positive Selection. Mol. Biol. Evol. 39, msab309 (2022).

99. Andrews, G. et al. Mammalian evolution of human cis-regulatory elements and transcription factor binding sites. Science 380, eabn7930 (2023).

100. Wang, K., Cai, B., Song, Y., Chen, Y. & Zhang, X. Somatosensory neuron types and their neural networks as revealed via single-cell transcriptomics. Trends Neurosci. 0, (2023).

101. Minnoye, L. et al. Chromatin accessibility profiling methods. Nat. Rev. Methods Primer 1, 1– 24 (2021).

102. Zeng, W. et al. Single-nucleus RNA-seq of differentiating human myoblasts reveals the extent of fate heterogeneity. Nucleic Acids Res. 44, e158 (2016).

103. Bakken, T. E. et al. Single-nucleus and single-cell transcriptomes compared in matched cortical cell types. PLoS ONE 13, e0209648 (2018).

104. Valtcheva, M. V. et al. Surgical extraction of human dorsal root ganglia from organ donors and preparation of primary sensory neuron cultures. Nat. Protoc. 11, 1877–1888 (2016).

105. Guimarães, M. Z. P. et al. Generation of iPSC-Derived Human Peripheral Sensory Neurons Releasing Substance P Elicited by TRPV1 Agonists. Front. Mol. Neurosci. 11, (2018).

106. Nickolls, A. R. et al. Transcriptional Programming of Human Mechanosensory Neuron Subtypes from Pluripotent Stem Cells. Cell Rep. 30, 932–946.e7 (2020).

107. Deng, T. et al. Scalable generation of sensory neurons from human pluripotent stem cells. Stem Cell Rep. 18, 1030–1047 (2023).

108. Lee, D. K. et al. Cloning and characterization of additional members of the G protein-coupled receptor family. Biochim. Biophys. Acta 1490, 311–323 (2000).

109. Bakken, T. E. et al. Comparative cellular analysis of motor cortex in human, marmoset and mouse. Nature 598, 111–119 (2021).

110. Zeng, H. What is a cell type and how to define it? Cell 185, 2739–2755 (2022).

111. Cunningham, F. et al. Ensembl 2022. Nucleic Acids Res. 50, D988–D995 (2022).

112. Ouyang, J. F., Kamaraj, U. S., Cao, E. Y. & Rackham, O. J. L. ShinyCell: simple and sharable visualization of single-cell gene expression data. Bioinformatics 37, 3374–3376 (2021).

113. Dimitrov, D. et al. Comparison of methods and resources for cell-cell communication inference from single-cell RNA-Seq data. Nat. Commun. 13, 3224 (2022).

114. Pawson, A. J. et al. The IUPHAR/BPS Guide to PHARMACOLOGY: an expert-driven knowledgebase of drug targets and their ligands. Nucleic Acids Res. 42, D1098–1106 (2014).

115. Shao, X. et al. CellTalkDB: a manually curated database of ligand-receptor interactions in humans and mice. Brief. Bioinform. 22, bbaa269 (2021).

116. Zeisel, A. et al. Cell types in the mouse cortex and hippocampus revealed by single-cell RNA-seq. Science 347, 1138–1142 (2015).

117. Yang, L., Tochitsky, I., Woolf, C. J. & Renthal, W. Isolation of Nuclei from Mouse Dorsal Root Ganglia for Single-nucleus Genomics. Bio-Protoc. 11, e4102 (2021).

118. Stuart, T. et al. Comprehensive Integration of Single-Cell Data. Cell 177, 1888–1902.e21 (2019).

119. Skene, N. G. et al. Genetic identification of brain cell types underlying schizophrenia. Nat. Genet. 1 (2018) doi:10.1038/s41588-018-0129-5.

120. Moretti, S. et al. Selectome update: quality control and computational improvements to a database of positive selection. Nucleic Acids Res. 42, D917–D921 (2014).

121. Hodge, R. D. et al. Conserved cell types with divergent features in human versus mouse cortex. Nature 573, 61–68 (2019).

122. Tran, M. N. et al. Single-nucleus transcriptome analysis reveals cell-type-specific molecular signatures across reward circuitry in the human brain. Neuron S0896-6273(21)00655–3 (2021) doi:10.1016/j.neuron.2021.09.001.

123. Hu, Y. et al. An integrative approach to ortholog prediction for disease-focused and other functional studies. BMC Bioinformatics 12, 357 (2011).

124. Khattak, S. et al. Germline Transgenic Methods for Tracking Cells and Testing Gene Function during Regeneration in the Axolotl. Stem Cell Rep. 1, 90–103 (2013).

125. Schloissnig, S., et al. The giant axolotl genome uncovers the evolution, scaling, and transcriptional control of complex gene loci. Proc. Natl. Acad. Sci. 118, e2017176118 (2021).

126. Lovely, A. M., Duerr, T. J., Stein, D. F., Mun, E. T. & Monaghan, J. R. Hybridization Chain Reaction Fluorescence In Situ Hybridization (HCR-FISH) in Ambystoma mexicanum Tissue. in Salamanders: Methods and Protocols (eds. Seifert, A. W. & Currie, J. D.) 109–122 (Springer US, 2023). doi:10.1007/978-1-0716-2659-7_6.

