## Supplementary material for "Harmonized cross-species cell atlases of trigeminal and dorsal root ganglia": Table S9 - Nomenclature across species

| Fiber Type | Atlas nomenclature | DRG Cre-lines | Cutaneous physiology | Mouse markers | Human markers | Macaque markers | Guinea Pig markers |
|---|---|---|---|---|---|---|---|
| A | Pvalb | Pvalb-cre[1] | Proprioceptors[1,2,3] | Pvalb | PVALB | PVALB | Pvalb |
| A | Ntrk3$^{high}$+S100a16 | Ntrk3-cre[4*] | Aβ SA1-LTMR / Aβ Field-LTMR[4] | Ntrk3$^{high}$+S100a16[a] / Ntrk3$^{high}$+Grm8 | NTRK3$^{high}$+GRM8 | NTRK3$^{high}$+S100A16[a] / NTRK3$^{high}$+GRM8 | Ntrk3$^{high}$+S100a16[a] |
| A | Ntrk3$^{high}$+Ntrk2 | Npy2r-cre[5*]/Ret-cre[6*] | Aβ-RA-LTMR[5,6] | Ntrk3$^{high}$+Ntrk2 / Ntrk3$^{high}$+Calb1[a] | NTRK3$^{high}$+NTRK2 | NTRK3$^{high}$+NTRK2 | † |
| A | Ntrk3$^{low}$+Ntrk2 | Ntrk2-cre[7*] | Aδ-LTMR[7] | Ntrk3$^{low}$+Ntrk2 | NTRK3$^{low}$+NTRK2 | NTRK3$^{low}$+NTRK2 | Ntrk3$^{low}$+Ntrk2 |
| A | Calca+Smr2 | Smr2-cre[8,9] | A-MH/C | Calca+Smr2[a] / Calca+Chrna7 | CALCA+CHRNA7 | CALCA+CHRNA7 | Tac1[b]+Chrna7 |
| A | Calca+Bmpr1b | Bmpr1b-cre[8,9] | A-M | Calca+Bmpr1b[a] | CALCA+KIT | CALCA+KIT | Tac1[b]+Kit |
| C | Th | Th-cre[10] | cLTMR[11] | Th[a] / Cdh9 | CDH9 | CDH9 | Cdh9 |
| C | Mrgprd | Mrgprd-cre[12,13] | C-MH[12,13] | Mrgprd[a] / Scn11a+Gfra1 | SCN11A+GFRA1 | SCN11A+GFRA1/ MRGPRD[a] | Scn11a+Gfra1 |
| C | Mrgpra3+Trpv1 | Mrgpra3-cre[*14,15,16] | C-MH[14,15,16] | Mrgpra3+Trpv1 | † | † | † |
| C | Mrgpra3+Mrgprb4 | Mrgprb4-cre[17,18,19] | C-MH[17,18,19] | Mrgpra3+Mrgprb4 | † | † | † |
| C | Calca+Sstr2 | Sstr2-cre[8,9] | C-H[8,9] | Calca+Sstr2[a] / Calca+Ptprt | CALCA+PTPRT | CALCA+PTPRT | † |
| C | Calca+Oprk1 | ? | ? | Calca+Oprk1 | † | † | † |
| C | Calca+Adra2a | Adra2a-cre[8,9] | ? | Calca+Adra2a | CALCA+ADRA2A | † | † |
| C | Calca+Dcn | ? | ? | Calca+Dcn | † | † | † |
| C | Rxfp1 | ? | ? | Rxfp1 | † | RXFP1 | † |
| C | Trpm8 | TrpM8-cre[19] /Trpm8-Flpo[9] | C-C[9,20,21,22] | Trpm8 | TRPM8 | TRPM8 | Trpm8 |
| C | Sst | Cysltr2-cre[9] / Sst-cre[23] | C-MH[9,23,24] | Sst | SST | SST | Sst |
| ? | Atf3 | Atf3-cre[25] | ? | Atf3 | ATF3 | ATF3 | Atf3 |

† Conserved marker genes could not be identified due to the low number of cells sequenced or absence in reference genome

[a] Cell type marker is specific in this species but nonspecific, unexpressed or absent in other species.

[b] Marker gene absent in reference genome
