## Supplementary material for "Harmonized cross-species cell atlases of trigeminal and dorsal root ganglia": Table S11 - Axolotl HCR probes

| Pool name | Sequence |
|---|---|
| TRPV1_B1 | gAggAgggCAgCAAACggAATGGCTGCCTCGCTGGCTCACTGCAG |
| TRPV1_B1 | CTCTCTTTCACGCTTCTCGACCAATTAgAAgAgTCTTCCTTTACg |
| TRPV1_B1 | gAggAgggCAgCAAACggAAAAAAGAACCTAAACATGAATCCCTT |
| TRPV1_B1 | CCTCATTCGTGCAAGTGGTGTCTCCTAgAAgAgTCTTCCTTTACg |
| TRPV1_B1 | gAggAgggCAgCAAACggAATCAATGGCAATGTGCAGCGCCGTCT |
| TRPV1_B1 | AGTTGCACCAGGTACATGTTCCTCCTAgAAgAgTCTTCCTTTACg |
| TRPV1_B1 | gAggAgggCAgCAAACggAATGGACGTCGGCCCCGTGCTGGATCA |
| TRPV1_B1 | CTGAAGAATTCTCCATCTGCCCTGGTAgAAgAgTCTTCCTTTACg |
| TRPV1_B1 | gAggAgggCAgCAAACggAATAAAAGCCTGGCTTTCCTTTAAGCT |
| TRPV1_B1 | GCAAGGGAGAGAGGAAGTTCACCAATAgAAgAgTCTTCCTTTACg |
| TRPV1_B1 | gAggAgggCAgCAAACggAAACGATGTCAATCTGGTTGGTGCAGG |
| TRPV1_B1 | TGGTACGGGTTCTGCAGGAGGTACTTAgAAgAgTCTTCCTTTACg |
| TRPV1_B1 | gAggAgggCAgCAAACggAAGAATCCTTGGTGGTGATGTTGGCCG |
| TRPV1_B1 | AAGGCATGAAAGACGGTGTTGCCTTTAgAAgAgTCTTCCTTTACg |
| TRPV1_B1 | gAggAgggCAgCAAACggAATCTTTGGTGTTGTCAGCAATGTCCA |
| TRPV1_B1 | TACATCTTGGTCACAAACTTTGTGTTAgAAgAgTCTTCCTTTACg |
| TRPV1_B1 | gAggAgggCAgCAAACggAATCAAGGTGGGGTTAATCTTTGCCCC |
| TRPV1_B1 | CTTTCTTGTTCGTAATCTCCTCCATTAgAAgAgTCTTCCTTTACg |
| TRPV1_B1 | gAggAgggCAgCAAACggAATTGCAGCAAGGGTTAAGGGAGTGAG |
| TRPV1_B1 | ATGCAAAGACCCCGATCTTGCCTGTTAgAAgAgTCTTCCTTTACg |
| TRPV1_B1 | gAggAgggCAgCAAACggAAGATCCTTGATCTCCCTCCGCAGTAT |
| TRPV1_B1 | ACTTGCGAGACAGATGCCTGCACTCTAgAAgAgTCTTCCTTTACg |
| TRPV1_B1 | gAggAgggCAgCAAACggAAGCACAGGTCCATAGGCCCACTCTGT |
| TRPV1_B1 | CACAAGACAGATCATAGAGGGACGATAgAAgAgTCTTCCTTTACg |
| TRPV1_B1 | gAggAgggCAgCAAACggAAGCACAGAGTTGTTCTCGTATGTGTC |
| TRPV1_B1 | TCTCACTACTGTAGGCGATGATCTCTAgAAgAgTCTTCCTTTACg |
| TRPV1_B1 | gAggAgggCAgCAAACggAACCAGCACCATGTCATGACGATTCGG |
| TRPV1_B1 | CCTGCAGCAACCGGTTGAGGGGTTCTAgAAgAgTCTTCCTTTACg |
| TRPV1_B1 | gAggAgggCAgCAAACggAATGCGCTTGACAAATCGGTCCCACTT |
| TRPV1_B1 | TGTACACTAGGAAGTTGAAGTAGAATAgAAgAgTCTTCCTTTACg |

| | |
|---|---|
| TRPV1_B1 | gAggAgggCAgCAAACggAACGATGGTGAAAACAATCATGTATGC |
| TRPV1_B1 | TTCCATCCACAGGCCTATAGTAGGCTAgAAgAgTCTTCCTTTACg |
| TRPV1_B1 | gAggAgggCAgCAAACggAATTGAATGCTCAATTGGGAAGGGAGG |
| TRPV1_B1 | CTCCAGCGTAGCGCAGATAAGCGTCTAgAAgAgTCTTCCTTTACg |
| TRPV1_B1 | gAggAgggCAgCAAACggAAAGATTCCTCCGATCACAGTGGTGAT |
| TRPV1_B1 | AATATTGAATCCCTCGGATGAAGAATAgAAgAgTCTTCCTTTACg |
| TRPV1_B1 | gAggAgggCAgCAAACggAATCTTCAGAGAAGGGCGCCTTTGCAC |
| TRPV1_B1 | CCTCACTGTAGCTGTCAATGAACAATAgAAgAgTCTTCCTTTACg |
| TRPV1_B1 | gAggAgggCAgCAAACggAAGGAACACCGACTGCACAAAGAACAG |
| TRPV1_B1 | TGAAGTACAGCACCACCGCCGTCAGTAgAAgAgTCTTCCTTTACg |
| TRPV1_B1 | gAggAgggCAgCAAACggAAGGGAGCCCACATACTCCTGCATGCC |
| TRPV1_B1 | CCCAACTCAGAGACAAGCACATGACTAgAAgAgTCTTCCTTTACg |
| TRPV1_B1 | gAggAgggCAgCAAACggAAATCCGCGAGTGTAGTACAGCATGTT |
| TRPV1_B1 | TGACTGAGTAGATGCCCATCACCTGTAgAAgAgTCTTCCTTTACg |
| TRPV1_B1 | gAggAgggCAgCAAACggAAAATCTCTCAGGATCATCTTCTCAAT |
| TRPV1_B1 | CACTGTAAACAAACATGAAGCGCAATAgAAgAgTCTTCCTTTACg |
| TRPV1_B1 | gAggAgggCAgCAAACggAAAAGCTGCAGCAAATCCAAAGAGGAA |
| TRPV1_B1 | ATTCGCCGTCCTCAATCAGGGTGACTAgAAgAgTCTTCCTTTACg |
| TRPV1_B1 | gAggAgggCAgCAAACggAAGCGACGGTGAGCTGCTTTCACCGAC |
| TRPV1_B1 | AGCTGGGAAAATAGCTTCCTTCATTTAgAAgAgTCTTCCTTTACg |
| TRPV1_B1 | gAggAgggCAgCAAACggAACCCCACGTCTTGTTTGGCCAGAGCC |
| TRPV1_B1 | TGTAGGACTCCGGACCACCTTTGCATAgAAgAgTCTTCCTTTACg |
| TRPV1_B1 | gAggAgggCAgCAAACggAAAGTTCTCAGTGAACTCCAAGTCCCC |
| TRPV1_B1 | GAAAGATGAAGATGGGCTTGAATCGTAgAAgAgTCTTCCTTTACg |
| TRPV1_B1 | gAggAgggCAgCAAACggAAATGTCATGATGACGTACACAACCAA |
| TRPV1_B1 | CAATGAGCATGTTGAGCAGCAGGATTAgAAgAgTCTTCCTTTACg |
| TRPV1_B1 | gAggAgggCAgCAAACggAATTTTGCTCACGGTCTCACCCATGAG |
| TRPV1_B1 | TCCAGATGCTCTTGCTTTCCTGCGCTAgAAgAgTCTTCCTTTACg |
| TRPM8_B2 | ACAGCTCAAGAACTTCTTCAGGTTCAAATCATCCAgTAAACCgCC |
| TRPM8_B2 | CCTCgTAAATCCTCATCAAACTCTGAATAAAGCTCAGTGAGCACG |

| | |
|---|---|
| TRPM8_B2 | GTTCCGGAACACCAGGGCGCTGAAGAAATCATCCAgTAAACCgCC |
| TRPM8_B2 | CCTCgTAAATCCTCATCAAAGTTGTATGAGTTCTTGGCGATGTGC |
| TRPM8_B2 | CTTCCACACGAAGGTGAGCAGTGTGAAATCATCCAgTAAACCgCC |
| TRPM8_B2 | CCTCgTAAATCCTCATCAAACTTGCTTCTCTGGAAGTTGGTGACC |
| TRPM8_B2 | GCCATCCTTCATGCTGTCATCTTCCAAATCATCCAgTAAACCgCC |
| TRPM8_B2 | CCTCgTAAATCCTCATCAAAGTAACGTCCTACTATTTGAACTTCC |
| TRPM8_B2 | GTGTCTTGTGATTGGCGATGCGTCAAAATCATCCAgTAAACCgCC |
| TRPM8_B2 | CCTCgTAAATCCTCATCAAATGCCCAGATGAAGAGGGCTTGTAGC |
| TRPM8_B2 | GGAGAGCTCCCTTTTGTTCTGCAGCAAATCATCCAgTAAACCgCC |
| TRPM8_B2 | CCTCgTAAATCCTCATCAAATCCTCTGGTCTGTTCCCAAATGACT |
| TRPM8_B2 | GCTAGCTCCCAGTGCAGCAAGTGTGAAATCATCCAgTAAACCgCC |
| TRPM8_B2 | CCTCgTAAATCCTCATCAAAGACTTTGGCCAAGCTCTTCAACAGC |
| TRPM8_B2 | CTCCCCAGCGGCATTGATATCATTCAAATCATCCAgTAAACCgCC |
| TRPM8_B2 | CCTCgTAAATCCTCATCAAACTCATACTCGTTGGCAATTTCCTCT |
| TRPM8_B2 | TTCGCTGAAGAGTTCGACTGCACGGAAATCATCCAgTAAACCgCC |
| TRPM8_B2 | CCTCgTAAATCCTCATCAAAAGCAAGGTCCTCGTCGCTGCTGTAG |
| TRPM8_B2 | TTCACAGGAATAGACCAGCAGCTGCAAATCATCCAgTAAACCgCC |
| TRPM8_B2 | CCTCgTAAATCCTCATCAAAGCCAGCTCGAGGCAGTTGCTGCCCC |
| TRPM8_B2 | ATAAACTGCTGATCTTTTGCTTCCAAAATCATCCAgTAAACCgCC |
| TRPM8_B2 | CCTCgTAAATCCTCATCAAAAGGAAGTTCTGGACTCCTGGCTGTG |
| TRPM8_B2 | GATATATGGCCATACCACTGCTTGGAAATCATCCAgTAAACCgCC |
| TRPM8_B2 | CCTCgTAAATCCTCATCAAAATGATCTTCCAGTTCTTTGTGTCAC |
| TRPM8_B2 | ATCAGCGGGAGGAAGAGGAGGCACAAAATCATCCAgTAAACCgCC |
| TRPM8_B2 | CCTCgTAAATCCTCATCAAATTCCTGAAGGAGATGAAGCCGCAGC |
| TRPM8_B2 | TGGGTCTGCTTTCCATCCAAGGGCTAAATCATCCAgTAAACCgCC |
| TRPM8_B2 | CCTCgTAAATCCTCATCAAAGTGATGAAGGCCCCGTACTTCCACA |
| TRPM8_B2 | GTCCAGGAGAAGACCACAAACGGTGAAATCATCCAgTAAACCgCC |
| TRPM8_B2 | CCTCgTAAATCCTCATCAAAAGGAGGAAGCCAATGTAGAAAATGG |
| TRPM8_B2 | TTCATCAGCAGCACGTAGGCGAAGAAAATCATCCAgTAAACCgCC |
| TRPM8_B2 | CCTCgTAAATCCTCATCAAATCTAGGCCAGTGGGCACGGGCTGGA |

| | |
|---|---|
| TRPM8_B2 | ACAAAGACCAGCCCATAGACAGAAAAAATCATCCAgTAAACCgCC |
| TRPM8_B2 | CCTCgTAAATCCTCATCAAAAACTGTCTGATTTCATCACAAAGGA |
| TRPM8_B2 | GTGAAATACTTGATACCGCTCATGTAAATCATCCAgTAAACCgCC |
| TRPM8_B2 | CCTCgTAAATCCTCATCAAAAGCATGTCCATGATGTTCCACATGT |
| TRPM8_B2 | ATGCCGGTGATGAAGTACAGGATGCAAATCATCCAgTAAACCgCC |
| TRPM8_B2 | CCTCgTAAATCCTCATCAAAGGGTTGGATTTATGTAGTCTGAACA |
| TRPM8_B2 | ATGACCCTCCCTGCGTACAGGGAGCAAATCATCCAgTAAACCgCC |
| TRPM8_B2 | CCTCgTAAATCCTCATCAAAGTGAAGATAATGTAGTCCAGGCAGA |
| TRPM8_B2 | ACAGTGAATATGTGGATCAGTCGGAAAATCATCCAgTAAACCgCC |
| TRPM8_B2 | CCTCgTAAATCCTCATCAAAATGATCTTGGGGCCCAGGTTCCGGC |
| TRPM8_B2 | ACATCGATTAACATTCTCTGCAGCAAAATCATCCAgTAAACCgCC |
| TRPM8_B2 | CCTCgTAAATCCTCATCAAAACCGCAAAGAGGAACAGGAAGAAGA |
| TRPM8_B2 | CGAGCCACGCCAAATGCAATCACCCAAATCATCCAgTAAACCgCC |
| TRPM8_B2 | CCTCgTAAATCCTCATCAAATGCTCGTTCAGCCTCAGGATGCCTT |
| TRPM8_B2 | ACAGAGCGGAAGATCCACTCCCAGCAAATCATCCAgTAAACCgCC |
| TRPM8_B2 | CCTCgTAAATCCTCATCAAAAACACGGCCAGATATGGCTCGTAGA |
| TRPM8_B2 | CCATCAACGTCGGAAGGGTACTGGCAAATCATCCAgTAAACCgCC |
| TRPM8_B2 | CCTCgTAAATCCTCATCAAAGTGCAGTGCTCGAAATCATAGGTGG |
| TRPM8_B2 | AGAGGTTTCGACTCATTCCCAGTGAAAATCATCCAgTAAACCgCC |
| TRPM8_B2 | CCTCgTAAATCCTCATCAAATGTTCGTCGGAATCCATCTCCACAC |
| TRPM8_B2 | ATGGTGATCCACTCTGGGAAGCGGGAAATCATCCAgTAAACCgCC |
| TRPM8_B2 | CCTCgTAAATCCTCATCAAAGACAGCATGTAGATGCACACAAGTG |
| TRPM8_B2 | AGGAGGTTGACCAGCAGGATGTTGGAAATCATCCAgTAAACCgCC |
| TRPM8_B2 | CCTCgTAAATCCTCATCAAACCAACGGTGTACCCAAACATGGCAA |
| TRPM8_B2 | ACCTGGTCATTGTTCTCTTGCACAGAAATCATCCAgTAAACCgCC |
| TRPM8_B2 | CCTCgTAAATCCTCATCAAAACCAGGAAGTAGCGCTGGAACTTCC |
| TRPM8_B2 | ATGGTGAGCCGGTTGCAGTACTCCTAAATCATCCAgTAAACCgCC |
| TRPM8_B2 | CCTCgTAAATCCTCATCAAATAGGCGAAGATCACAAACGGGAATG |
| TRPM8_B2 | AGCATCTTCCTCAGCACCATGAAGAAAATCATCCAgTAAACCgCC |
| TRPM8_B2 | CCTCgTAAATCCTCATCAAAGGGGCCTCCTCCTTCCAGCAGCAGT |

| | |
|---|---|
| TRPM8_B2 | TTTCTCGAACAGCATACGGTGGGCTAAATCATCCAgTAAACCgCC |
| TRPM8_B2 | CCTCgTAAATCCTCATCAAATCCCAGGCCAGGGTCTCTGTGTCCT |
| TRPM8_B2 | ATGAGATAGTTCTCTTTCATGACAGAAATCATCCAgTAAACCgCC |
| TRPM8_B2 | CCTCgTAAATCCTCATCAAAGAGTCATTGGCCTTGATGTTGCTTT |
| TRPM8_B2 | CGGAACCGGTGCTCCATCTGCTCCGAAATCATCCAgTAAACCgCC |
| TRPM8_B2 | CCTCgTAAATCCTCATCAAAAGGTCGTGCAGCTTGCTCTCCATCT |
| TRPM8_B2 | CTGGTAATCTCTTTCATGATGCGCTAAATCATCCAgTAAACCgCC |
| TRPA1_B3 | gTCCCTgCCTCTATATCTTTGTAGGATTTAATAGTAATTTAATAC |
| TRPA1_B3 | TTGTGTTGAGGCAAAGATCCAGACCTTCCACTCAACTTTAACCCg |
| TRPA1_B3 | gTCCCTgCCTCTATATCTTTCTATTTGTCCAGCGGCATCTCTTTG |
| TRPA1_B3 | TTGCCTTCTCAGCTGCCCCTAACACTTCCACTCAACTTTAACCCg |
| TRPA1_B3 | gTCCCTgCCTCTATATCTTTTGACTTATGCTATCACTGCAGTCAG |
| TRPA1_B3 | AGTAGCACAGTCAGGCAATCAGGGATTCCACTCAACTTTAACCCg |
| TRPA1_B3 | gTCCCTgCCTCTATATCTTTCCGACGGCCAAACCAATCTCCACAG |
| TRPA1_B3 | GCATTGCGTTGTACTTCTGCAATGTTTCCACTCAACTTTAACCCg |
| TRPA1_B3 | gTCCCTgCCTCTATATCTTTACCTGCATTTCAATTCTTTTAAGTG |
| TRPA1_B3 | TTCTTTTCCAAGCTGGTGTGAAGGTTTCCACTCAACTTTAACCCg |
| TRPA1_B3 | gTCCCTgCCTCTATATCTTTACTCGTTTTAGCAACCAATAAGGCA |
| TRPA1_B3 | GGATAGACAGTTATGGAGGTCTTATTTCCACTCAACTTTAACCCg |
| TRPA1_B3 | gTCCCTgCCTCTATATCTTTGCAACTCCCCAGATTCTTGGCCTGT |
| TRPA1_B3 | ATGAACAGGAGCAAAGGTGAATTGATTCCACTCAACTTTAACCCg |
| TRPA1_B3 | gTCCCTgCCTCTATATCTTTTCTGGTGTTGATCCCTCACAAGCAA |
| TRPA1_B3 | TCCACAGCAGTGTCGGTAGGCGGCATTCCACTCAACTTTAACCCg |
| TRPA1_B3 | gTCCCTgCCTCTATATCTTTCTGTGCTTTTGCTTTAAAAGTTCCG |
| TRPA1_B3 | TGAAGCAGAGAAGACATGTCTTTCATTCCACTCAACTTTAACCCg |
| TRPA1_B3 | gTCCCTgCCTCTATATCTTTATCAGTTTAATTAGTTCATGTTGTT |
| TRPA1_B3 | TCTGAAATAATTTCCATTTTCTGAATTCCACTCAACTTTAACCCg |
| TRPA1_B3 | gTCCCTgCCTCTATATCTTTTGATCCTGCTCCTCCTCATCTTCAG |
| TRPA1_B3 | TTTTGACTTTTGACCTTGCAACTTTTTCCACTCAACTTTAACCCg |
| TRPA1_B3 | gTCCCTgCCTCTATATCTTTTTCCATTTGCTGTCCTTGCGGTTCA |

| | |
|---|---|
| TRPA1_B3 | TTTGCTTTGACTGCTTTTAGCACACTTCCACTCAACTTTAACCCg |
| TRPA1_B3 | gTCCCTgCCTCTATATCTTTAACAGTGAGTCTCTGGTTCACACAG |
| TRPA1_B3 | CAATATCTGATTATACTATCAAACATTCCACTCAACTTTAACCCg |
